# Aromatic natural products synthesis from aromatic lignin monomers using *Acinetobacter baylyi* ADP1

**DOI:** 10.1101/2023.08.24.554694

**Authors:** Bradley W. Biggs, Keith E. J. Tyo

## Abstract

Achieving sustainable chemical synthesis and a circular economy will require process innovation to minimize or recover existing waste streams. Valorization of lignin biomass has the ability to advance this goal. While lignin has proved a recalcitrant feedstock for upgrading, biological approaches can leverage native microbial metabolism to simplify complex and heterogeneous feedstocks to tractable starting points for biochemical upgrading. Recently, we demonstrated that one microbe with lignin relevant metabolism, *Acinetobacter baylyi* ADP1, is both highly engineerable and capable of undergoing rapid design-build-test-learn cycles, making it an ideal candidate for these applications. Here, we utilize these genetic traits and ADP1’s native β-ketoadipate metabolism to convert mock alkali pretreated liquor lignin (APL) to two valuable natural products, vanillin-glucoside and resveratrol. En route, we create strains with up to 22 genetic modifications, including up to 8 heterologously expressed enzymes. Our approach takes advantage of preexisting aromatic species in APL (vanillate, ferulate, and *p*-coumarate) to create shortened biochemical routes to end products. Together, this work demonstrates ADP1’s potential as a platform for upgrading lignin waste streams and highlights the potential for biosynthetic methods to maximize the existing chemical potential of lignin aromatic monomers.

## Introduction

Progress towards sustainable chemical synthesis will require a transition away from the current linear model of manufacturing and consumption, where resources are extracted, fashioned, utilized, and discarded, to one that is instead circular^1^. Non-food biomass has significant potential as a feedstock for sustainable chemical synthesis because of its inherent linkage to circularity through photosynthetic CO_2_ fixation and because biomass utilization, in exchange of non-renewable feedstocks, can alleviate ecosystem burden^2,3^. To be successful, though, biomass-based processes must be efficient and be capable of generating a broad array of products, including high value products, especially in view of competing traditional processes^4,5^.

A present and acute need within biomass-based processes is for more complete biomass utilization^6,7^. Specifically, lignin utilization has lagged behind that of the other two primary components (cellulose and hemicellulose)^8^. Lignin, which provides the structure for plants, is a heterogenous aromatic polymer, and its complex nature has proved challenging for traditional means of upgrading^9^. Instead, lignin is often treated as a waste product of biomass processes and is burned for heat^10^. Creating new processes that generate value-added products from lignin would represent a significant advance for carbon circularity broadly and biomass circularity in particular.

Microbial systems have been proposed as a solution for lignin valorization, as part of a “biological funneling” concept^11^ (Figure 1). In this approach, lignin is pretreated to generate a mixture of aromatic monomers, such as *p*-coumarate, ferulate, vanillate, and *p*-hydroxybenzoate. Following, this mixture is fed to microbial systems that leverage the β-ketoadipate pathway to “funnel” the various lignin aromatic monomers to tractable starting points for biochemical upgrading. Previous work within the lignin biological funneling paradigm has primarily focused on selectively retaining a single metabolite (i.e. vanillin)^12^, converting all metabolites to commodity chemicals like muconic acid^13^, or building a pathway from the TCA cycle^14^, the entry point of the β-ketoadipate pathway to central carbon metabolism. Few, if any, approaches leverage the existing chemical potential of the available aromatic species. Utilizing this chemical potential to synthesize high value aromatic products from lignin could significantly impact the economic feasibility of lignin-based biomanufacturing and thus biomass utilization as a whole.

**Figure 1.**
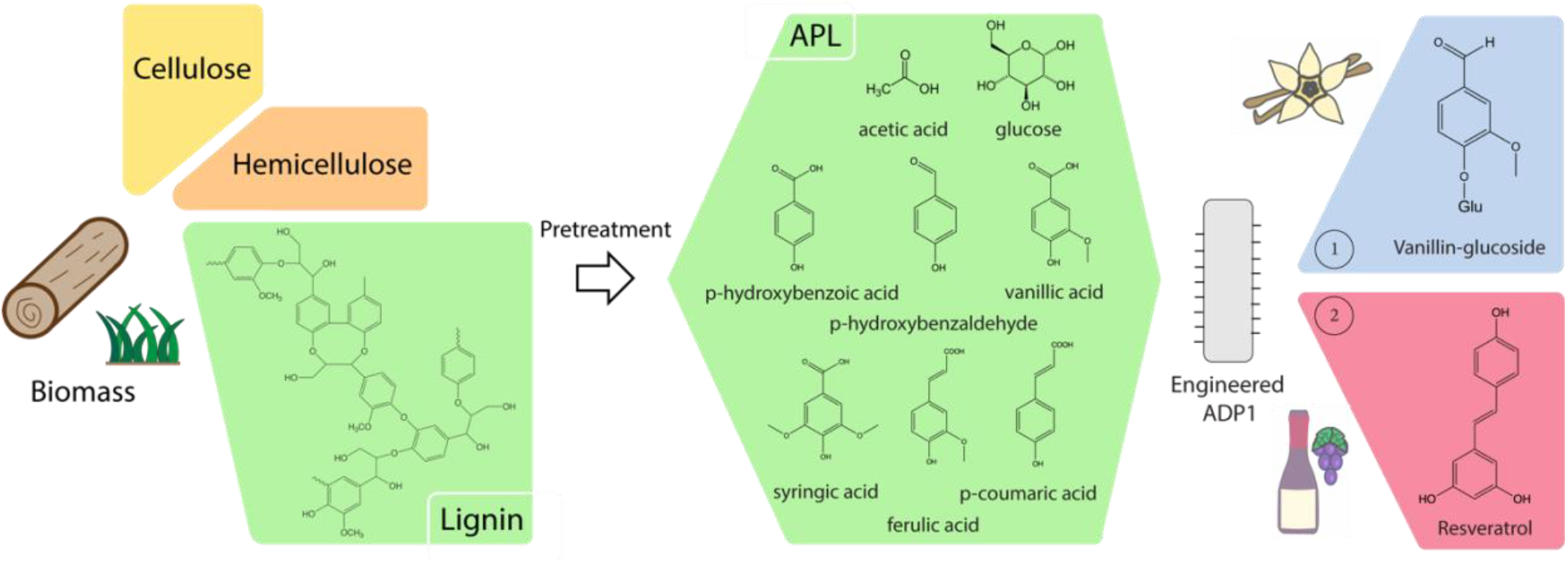
Lignin funneling for synthesis of natural products with ADP1. Figure shows the concept of taking a given lignin source, utilizing a chemical pretreatment step to convert it into individual aromatic monomers, and then feeding this carbon stream to a microbial host. At the stage of microbial feeding, the β-ketoadipate pathway is used to funnel aromatic lignin monomers into central carbon metabolism, providing starting points for metabolic engineering approaches. In this work the aromatic products vanillin-glucoside and resveratrol were produced.

Here, we use the bacterium *Acinetobacter baylyi* ADP1 to synthesize the aromatic natural products vanillin-glucoside and resveratrol from the existing aromatic monomer species of a mock alkali pretreated liquor lignin (APL)^15–17^ (Figure 1). Conscientious of long-term economic considerations, all processes in this work utilized this feedstock and in the context of a minimal medium (M9) with only trace metals supplemented. Vanillin-glucoside and resveratrol were chosen because, as with many valuable natural products, their demand outpaces natural sourcing, where low native yields and arable land requirements constrain scalability^18–20^. Motivated by this constraint and consumer preferences away from petroleum-derived products, metabolic engineering approaches have previously been developed as a sustainable solution for scaled production of each of these products, primarily using glucose as a feedstock^21,22^. By instead synthesizing these products from lignin aromatics, we take advantage of significantly shorter biosynthetic pathways, simultaneously leveraging ADP1’s native metabolism of ferulate (vanillin-glucoside) and *p*-coumarate (ferulate).

ADP1 was chosen as a host organism because of its possession of the β-ketoadipate pathway for aromatic monomer catabolism and, more importantly, its engineerability that we have previously demonstrated^23^. Biomanufacturing success is largely a function of the ability to modify the host organism toward engineering goals^24^. ADP1 excels in this area, and its facile genetics enabled the creation of strains with up to 22 distinct scarless genetic knock outs and 8 heterologously expressed enzymes. Taken together, our works is a key proof-of-concept demonstration for high-value aromatic product synthesis from waste lignin biomass.

## Results and Discussion

We begin with our biological funneling approach for vanillin-glucoside production from mock APL (Figure 2). Like the latter steps of glucose-based approaches to vanillin or vanillin-glucoside synthesis^22,25–28^, our synthesis pathway proceeds from protocatechuate (PCA) to vanillate by catechol O-methyltransferase (COMT), from vanillate to vanillin by carboxylic acid reductase (Car), and from vanillin to vanillin-glucoside by a UDP-glucose dependent glycosyltransferase (UGT) (Figure 2). Vanillin-glucoside was prioritized as a final product over vanillin because it has previously been shown less toxic to production hosts, because of its superior aqueous solubility, and as it naturally exists in *Vanilla planifolia*^22,26,29^.

**Figure 2.**
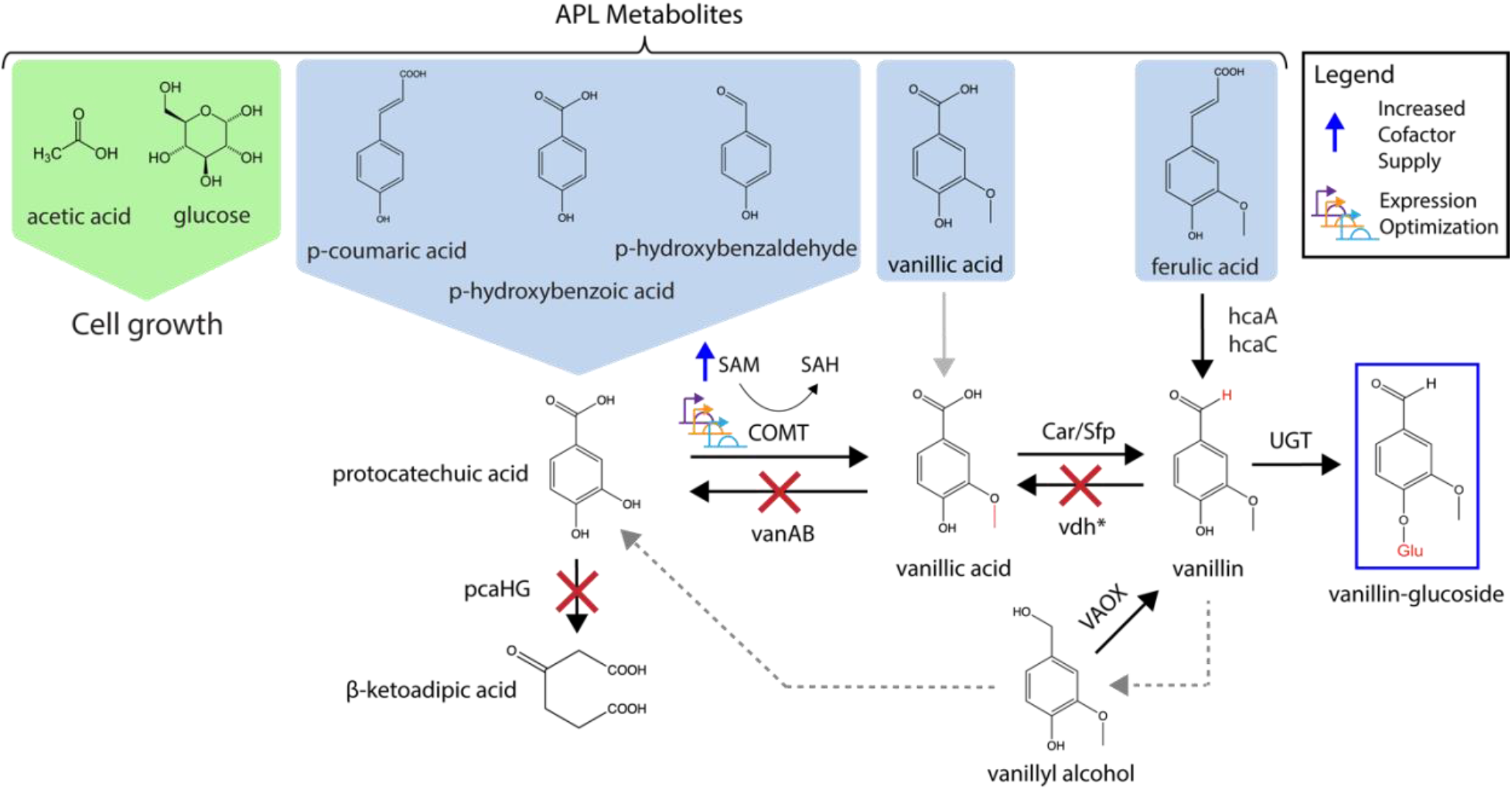
Vanillin-glucoside synthesis strategy from mock APL. Figure depicts the overall approach, wherein mock APL components (top section in the green and light blue shaded areas) are funneled into a metabolic engineering pathway towards vanillin-glucoside. Three genes, a catechol O-methyltransferase (COMT), carboxylic acid reductase (Car), and UDP-glucose dependent glycosyltransferase (UGT) are used to actively synthesize vanillin-glucoside from lignin aromatics (blue shading), while glucose and acetate (green shading) are devoted to cell growth. Reverse reactions, including a promiscuous dehydrogenase side degradation of vanillin to vanillyl alcohol that can be reconverted to vanillin by vanillyl alcohol oxidase (VAOX), discussed in detail below, are also shown. Red “Xs” are shown over reverse reactions that have been removed from ADP1, with the gene removed listed except for the case of vdh*, which represents many enzymes. Lastly, engineering strategies to improve the first step of the pathway (COMT) are indicated (co-factor supply increase and expression optimization).

Our biosynthetic pathway begins with PCA owing to the composition of our feedstock (APL) and ADP1’s metabolism of it. Mock APL, which based on an alkali pretreated corn stover^15–17^, contains glucose, acetate, and the aromatic monomers *p*-coumarate, *p*-hydroxybenzoate, *p*-hydroxybenzaldehyde, vanillate, and ferulate. As mentioned, ADP1 metabolizes the aromatic monomers of mock APL via the β-ketoadipate pathway, which funnels these aromatics to either catechol or PCA^30^. Importantly, all of the aromatic monomers in APL belong to the PCA branch of the β-ketoadipate pathway, and are thus funneled to PCA to form the basis for vanillin-glucoside synthesis. The remaining glucose and acetate of APL is used for cell growth and maintenance. It is also worth noting that one APL species (vanillate) is already part of the vanillin-glucoside synthesis pathway and that the first step of ferulate’s native degradation is to another a pathway metabolite (vanillin), allowing maximum utilization of the feedstock.

While fundamentally enabling for this study, ADP1’s aromatic funneling also presents a challenge. The capability to degrade various aromatic species to tractable starting points for metabolic engineering necessitates that vanillin-glucoside intermediate pathway metabolites can be metabolized unproductively for growth. To ensure that synthesis the pathway proceeds primarily the forward direction, and that all aromatic species can be utilized productively, we engineered ADP1 to inhibit or prevent PCA, vanillate, and vanillin consumption. First, the enzymes responsible for PCA and vanillate degradation in ADP1 were removed, *pcaHG* and *vanAB* respectively. A strain with both pairs of enzymes removed was fed 1 mM p-coumarate and 1 mM vanillin, along with glucose and acetate for growth. As expected, this strain retained PCA (derived from *p*-coumarate) and vanillate (derived from vanillin) (Figure 3).

**Figure 3.**
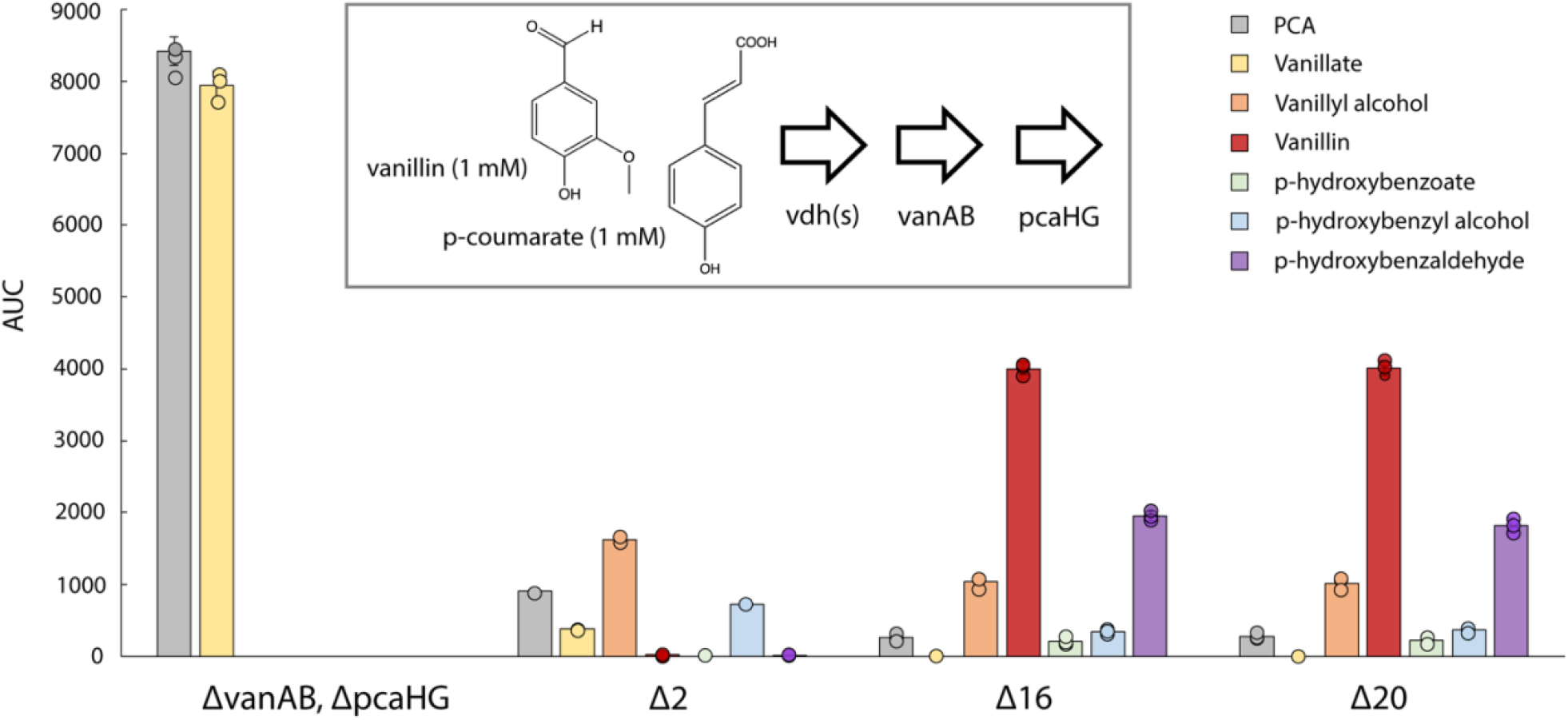
Comparison of strains with vanillin pathway metabolite degradation enzyme knock outs. Feeding 1 mM p-coumarate and 1 mM vanillin to ADP1 deletion strains. Figure shows the carbon consumption pattern of ADP1 strains with different gene deletions. All four strains have both *vanAB* and *pcaHG* deleted, with ΔvanAB, ΔpcaHG having only these four genes removed. As expected, by removing the genes capable of PCA (*pcaHG*) and vanillate (*vanAB*) degradation, metabolites are retained at PCA and vanillate. After the deletion of two putative vanillin dehydrogenase genes (*hcaB* and *areC*), strain Δ2 shows the accumulation of other compounds including vanillyl alcohol and *p*-hydroxybenzyl alcohol. After a series of additional putative vanillin dehydrogenases deletions, strains Δ16 and Δ20 showed similar degradation profiles, and measurable vanillin was retained. All error bars represent standard deviation for biological triplicate data, expect for Δ2, which only had biological replicate.

While removing the genes responsible for vanillate and PCA degradation was straightforward, removing the dehydrogenase activity responsible for vanillin degradation activity was not. While other microbes that possess the β-ketoadipate pathway have annotated vanillin dehydrogenase (*vdh*) enzymes, ADP1 lacks an enzyme with this specific annotation.

Therefore, we utilized Psi-BLAST^31^ searches of ADP1’s genome with known vanillin dehydrogenases from both *Pseudomonas* and *Sphingomonas* species and *ald* from *Corynebacterium glutamicum* to identify candidates for deletion (Tables S1). The top most likely candidates based on homology from this search were *hcaB* (annotated as hydroxybenzaldehyde dehydrogenase) and *areC* (annotated as benzaldehyde dehydrogenase II).

Both *hcaB* and *areC* were removed from the strain with *vanAB* and *pcaHG* already removed to create the strain Δ2, with Δ2 a reference to the number of putative vanillin dehydrogenase genes removed. When Δ2 was grown on the previously described medium with *p*-coumarate and vanillin, the retained product profile changed measurably (Figure 3). While some PCA and vanillate (the previously observed oxidation products) still appeared, new species emerged including vanillyl alcohol and *p*-hydroxybenzyl alcohol (reduction products). However, no vanillin was observed after the 24-hour cultivation, suggesting that alternative degradation pathways were being utilized by ADP1. Others have observed a similar need to remove multiple aldehyde dehydrogenase enzymes to prevent vanillin degradation^27,32^. Accordingly, we systematically deleted putative vanillin dehydrogenase enzymes from our candidate list prioritizing those with high homology to known vanillin dehydrogenases and with increased expression from growth on quinate (an aromatic molecule) compared to succinate (non-aromatic molecule) as determined by a prior study^33^ (Table S2 for list of genes removed).

At approximately strain Δ16, we identified a strain that retained vanillin along with *p*-hydroxybenzaldehyde, which is likely degraded by the same promiscuous dehydrogenases. Additional knocks outs neither appeared to improve vanillin retention nor change the retained product profile (Figure 3, comparing Δ16 and Δ20, Figures S1 for full set of knockouts up to Δ16). In addition, strains with more knockouts generally showed reduced growth, but only in the minimal medium condition with APL and not in LB (Figures S1-2). Therefore, Δ16 was used for subsequent work, as it enabled vanillin-glucoside pathway intermediate retention and thus provided a foundation for vanillin-glucoside pathway engineering in ADP1.

### Introduction of vanillin synthesis pathway to ADP1

Having obtained a strain capable of improved vanillin retention, individual enzymes in the vanillin-glucoside synthesis pathway were tested for activity in ADP1 with two studies used as the basis for the enzymes chosen for testing^22,27^. First, catechol O-methyltransferase (COMT) from *Homo sapiens* was tested and found capable of converting PCA to vanillin in ADP1 (Figure S3). Next, the UDP-glucose dependent glycosyl transferase (UGT) step that converts vanillin to vanillin-glucoside was tested. The glycosylation was tested second as ADP1 maintained some vanillin degradation capacity but is unable to degrade vanillin-glucoside, so any successful conversion of vanillin to vanillin-glucoside should be observable. For this UGT step, UGT72E2 of *Arabidopsis thaliana* was chosen and found capable of converting vanillin to vanillin-glucoside in ADP1 (Figure S4).

Finally, carboxylic acid reductase (Car) from *Nocardia iowensis* was tested for its ability to convert vanillate to vanillin. Car was paired both with its native phosphopantetheinyl transfer partner *ppt* and an alternative partner from *Bacillus subtilis* (*sfp*), which had been found successful in a previous work^27^. Car enzymatic tests were carried out in the context of a strain also expressing UGT to trap any vanillate converted to vanillin at vanillin-glucoside. While both pairings were successful, Car/sfp was the superior partnership, as it showed both greater amounts of vanillin-glucoside and vanillyl alcohol (a re-metabolization of vanillin) and as the full amount of vanillate was utilized (Figure S5). All three enzymes were expressed in the context of Δ16 with COMT integrated at *vanAB* under Trc expression, UGT integrated at *pcaHG* under Trc-BCD9 expression, and with Car/Sfp expressed via plasmid. In a 24-hour cultivation in M9 with simplified mock APL, vanillin-glucoside production was observed (Figure S6-7).

### Enhancing o-methyl transferase activity for PCA

Both our own testing (Figure S7) and prior work^25,28,34^ indicate an apparent bottleneck at the first step of the pathway with the conversion of PCA to vanillate. As all the aromatic metabolites present in mock APL are on the PCA side of the β-ketoadipate pathway, this could present an issue for fed-batch experiments where PCA could accumulate and become toxic. Not only is metabolite accumulation generally detrimental in metabolic engineering, but PCA is known to behave as an iron chelator^35^, which could cause further issues. Accordingly, we prioritized improving the conversion of PCA to vanillate.

Prior studies in *Escherichia coli* showed that conversion of PCA to vanillate can be limited by enzyme expression and cofactor availability^27^. Therefore, for an initial experiment, we tested increasing COMT expression using a plasmid, providing the precursor for the methyl donating cofactor s-adenosylmethionine (SAM) that COMT utilizes to convert PCA to vanillate (L-methionine), along with testing the addition of trace metals that have been shown to improve strain performance. Each intervention showed slight improvement for conversion of PCA to vanillate (Figure S8), providing direction for subsequent optimization.

To optimize COMT expression, we first modified the expression of the integrated COMT by testing different ribosomal binding site (RBS) variants to replace the initial “agga” RBS. For this we used bicistronic design (BCD) constructs^36^ that we had previously validated in ADP1^23^. Increased expression provided modest benefit up to a point, with BCD20 providing a 24% increase in turnover compared to the “agga” RBS. However, the constructs with higher expression strength (BCD14 and BCD9) showed reduced COMT turnover (Figure S9).

Interestingly, the high expression strength constructs also showed a sharp reduction in growth (Figure S9). As SAM is required for essential processes in the cell (phospholipid biosynthesis, protein post-translational modification, DNA methylation)^37^, this reduction could be the result of an exhaustion of SAM availability in the cell, which would make improvement of SAM availability critical for pathway performance.

Motivated by this possibility, we next sought to test the impact of modifying SAM availability. Leveraging previous work^28^ and the metabolic mapping of ADP1^38^, we identified six genes from *E. coli* that could be incorporated to potentially improve the SAM pool in ADP1, *mtn, luxS, metK, CysE, metA*, and *metB*. Figure S10 provides a “map” of SAM related enzymes, considering both ADP1 and *E. coli* pathways, that could help generate (or regenerate) the cofactor. Worth noting, ADP1 and *E. coli* have different homoserine to homocysteine pathways, with ADP1 preferring acetylation and *E. coli* preferring succinylation. In addition, by utilizing heterologous *E. coli* enzymes in ADP1, we hypothesized that we might avoid endogenous regulation.

While we observed a benefit with L-methionine addition (Figure S8), suggesting that the activity of metK may not be limiting, as was the finding of an *E. coli* study^28^. However, for thoroughness, we still tested the addition of metK and two enzymes involved in the recycling of SAH (the metabolic product of SAM after methyl donation), mtn and luxS. The six *E. coli* SAM genes were integrated in different groupings and altogether in the context of the Δ16 strain. Each SAM cofactor enzyme underwent a scarless and markerless integration and was integrated with a Trc promoter and lacO operator. In terms of integration location, metK was integrated at ACIAD2929, the feedback resistant mutant metA* at acoD, metB at ACIAD1577-ACIAD1578, the feedback resistant cysE* at dhbA, mtn at quiA, and luxS at calB.

Though several combinations showed benefit over the reference strain with no additional SAM enzyme expression, metK improved turnover the most (>4-fold improvement in COMT activity, Figure S11). The flux through this enzymatic step was increased to the point that a potential isomer “isovanillate” was observed (Figure S12), where a “shoulder” in the HPLC trace on the vanillate peak potentially represents a promiscuous para O-methylation in place of the desired meta O-methylation^34^. The best overall combination of SAM enzymes was that of luxS, mtn, and metK (given the shorthand name “lmmK”). Though on their own they provided benefit over the reference strain, the inclusion of CysE* and metA* with “lmmK” hindered turnover (Figure S11).

Combining the COMT expression and SAM pool optimization, a new strain was constructed from the Δ16 background with COMT under Trc-BCD20 chromosomal expression, the set of “lmmK” SAM enzymes chromosomally integrated, and with UGT chromosomally expressed as before (Trc-BCD9). Curiously, though this new strain showed a 48% improvement over the increased expression of COMT with BCD20 alone, and a 2.58-fold improvement over the reference strain of Δ16, it showed a reduction in activity compared to “lmmk” alone (Figure S13). This result, including the lower OD_600_ of this strain compared to “lmmk,” suggests that either a potential total capacity for heterologous expression has been reached or that the SAM pool is still being exhausted. In the context of the full vanillin-glucoside pathway, this strain showed a slight decrease in vanillin-glucoside titer from mock APL compared to the Δ16 reference strain giving 10.9 mg/L compared to 12.7 mg/L in a 24-hour cultivation in M9 with trace metals at the 3 mL scale (Figure S14). However, the strain did show a modest (8%) increase in vanillate compared to the reference strain (p-value 0.074) (Figure S15).

### Vanillyl alcohol recovery with vanillyl alcohol oxidase

During cultivations with the vanillin dehydrogenase knockout strains, vanillin degradation was still observed, even though the primary degradation pathway (from vanillin to vanillate) had been removed. One of the alternative vanillin degradation pathways involves the reduction of vanillin to vanillyl alcohol, with *p*-hydroxybenzaldehyde being similarly degraded to benzoyl alcohol instead of *p*-hydroxybenzoate (Figure S16). Both of these secondary degradation products were observed by HPLC in strains with greater numbers of putative vanillin dehydrogenases removed (Figure 3). As it would be desirable to recapture this carbon for vanillin-glucoside production, we tested an enzyme known as vanillyl alcohol oxidase (VAOX)^39,40^, which had been previously used in an enzyme cascade to synthesize vanillin^41^, for activity in ADP1. Feeding vanillyl alcohol to ADP1, we observe VAOX activity (Figure S17), indicating that this enzyme could be useful to include in our engineered strain. Based on the results of the COMT optimization in the context of the full pathway, we integrated VAOX into a strain with only metK expressed among SAM cofactor enzymes and COMT expressed under a weak RBS. This strain showed 17.1 mg/L vanillin-glucoside, along with 19.5 mg/L vanillin, for a total of 36.6 mg/L of the two primarily flavor compounds (Figure 4).

**Figure 4.**
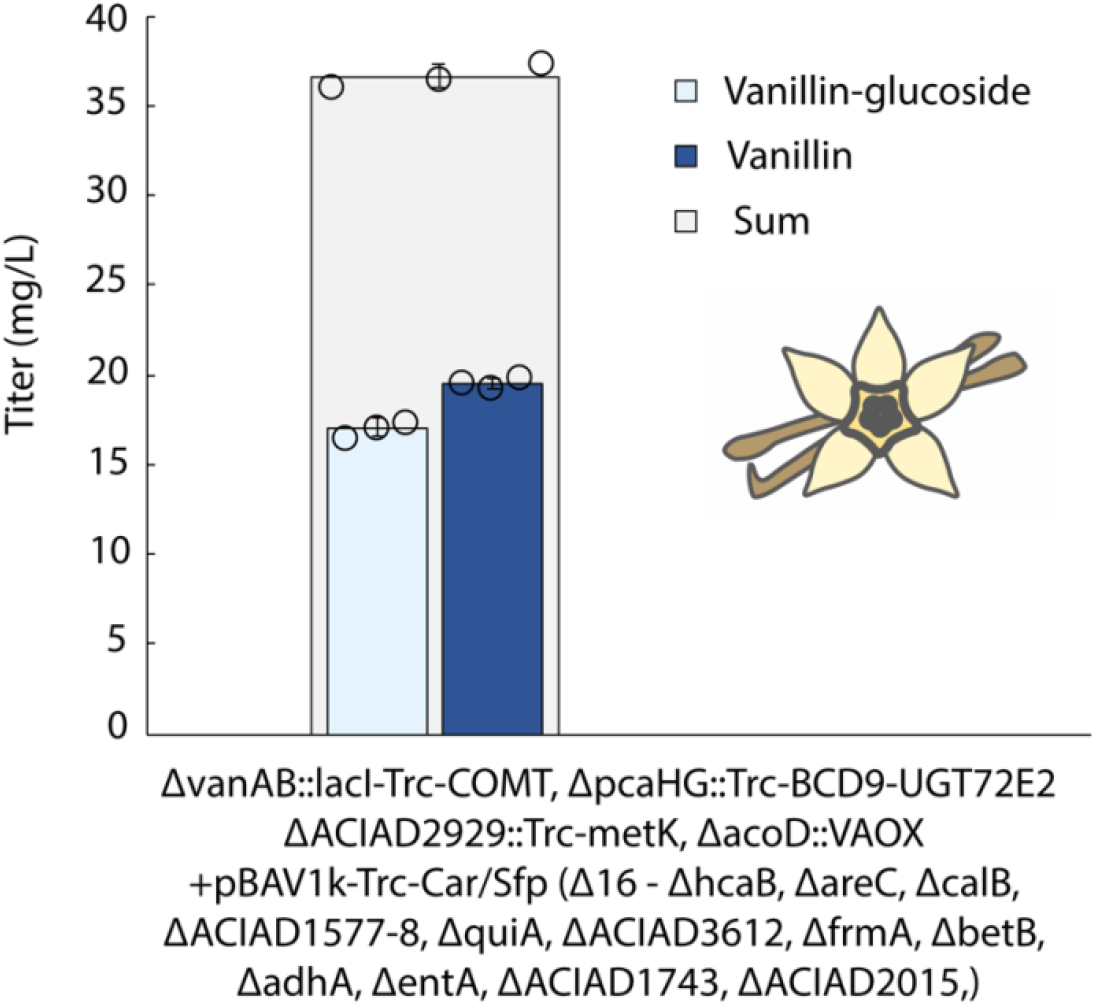
Vanillin pathway production. Titers for the cultivation of the full vanillin-glucoside pathway with metK and VAOX additionally expressed.

### Resveratrol production

In an effort to demonstrate that this approach with ADP1 has applicability as platform for lignin upgrading beyond vanillin-glucoside synthesis, we sought to identify an additional natural product that could be made from aromatic lignin monomers. Because the highest concentration lignin monomer in APL is *p*-coumarate, we wanted to target a product that could be readily made from this aromatic species. To this end, *p*-coumarate is only two biosynthetic steps away from the valuable nutraceutical flavonoid resveratrol (Figure 5, top), which has been a frequent target of metabolic engineers due to limits for plant extraction (grape) and chemical synthesis^21^.

**Figure 5.**
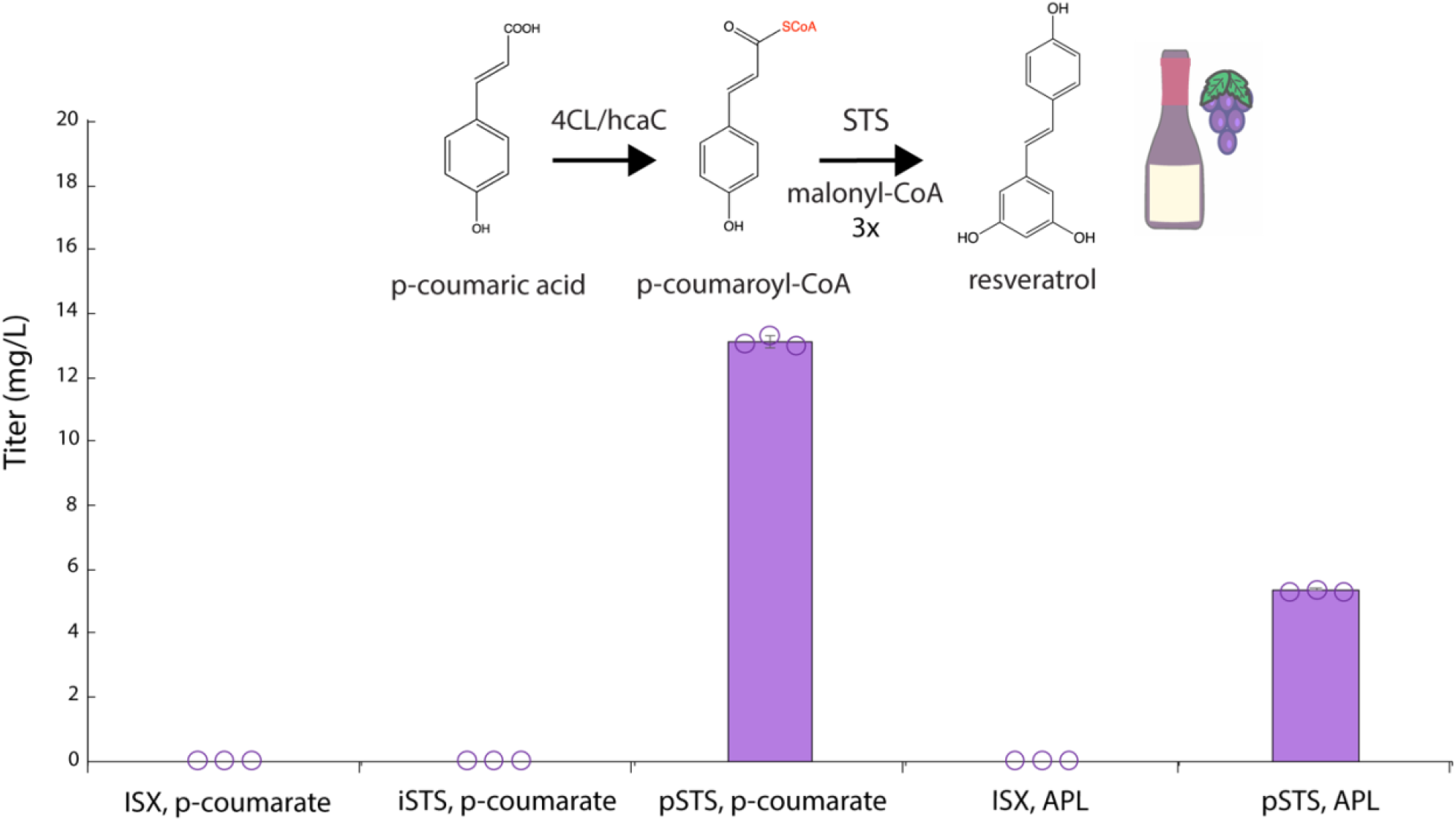
Resveratrol production in ADP1. Top of the figure shows the resveratrol pathay from p-coumaric acid. Below the graph shows resveratrol production from ADP1 expression of resveratrol synthase (STS) either chromosomally (iSTS) or by plasmid (pSTS) for medium conditions either containing *p*-coumarate as the sole aromatic monomer or full mock APL.

To date, approaches for resveratrol biosynthesis have been primarily carried out in common metabolic chassis strains (*E. coli, S. cerevisiae*) using traditional feedstocks such as glycerol and sucrose, often supplementing L-tyrosine or *p*-coumarate to the medium^42,43^. These approaches leverage amino acid biosynthesis and subsequently convert tyrosine to *p*-coumarate by tyrosine ammonia lyase (TAL)^44^. Following, *p*-coumarate is converted to resveratrol by two additional enzymatic steps^45^. First, 4-coumarate-CoA ligase (4CL) is used for CoA addition^44^. Second, resveratrol synthase (stilbene synthase, STS) converts *p*-coumaroyl-CoA to resveratrol using three units of malonyl-CoA^45^. Though a simple pathway, resveratrol synthesis is intriguing because of its inherent value, as it can function as a proof of concept for the broader flavonoid family of molecules^46^, and because resveratrol can form the basis for further decoration to create derivatives such as the methylated pinostilbene or pterostilbene^47^.

Our approach to resveratrol synthesis is shown in Figure S18. Fortuitously, the first step of resveratrol biosynthesis from *p*-coumarate and ADP1’s native metabolism of *p*-coumarate and ferulate are the same CoA ligation (Figure S19). To determine if the activity of the enzyme often used for resveratrol synthesis (4CL of *Arabidopsis thaliana*) was interchangeable with ADP1’s native enzyme for this activity (hcaC), we first deleted *hcaC*. As expected, a Δ*hcaC* strain of ADP1 was not able to degrade *p*-coumarate or ferulate (Figure S20). Next, to test if 4CL was interchangeable for hcaC, we added 4CL under plasmid expression (pBAV1k-lacI-Trc-4CL) into Δ*hcaC*. Complementing the Δ*hcaC* strain with 4CL restored degradation of *p*-coumarate and ferulate (Figure S20), suggesting that ADP1 could perform the first step of resveratrol synthesis from *p*-coumarate without heterologous expression of 4CL.

The interchangeability of hcaC and 4CL for CoA-ligation of *p*-coumarate and ferulate provides both an advantage and a disadvantage. Helpfully, it reduces the number of heterologous enzymes necessary to synthesize resveratrol in ADP1. However, it prevents the metabolite pooling approach used for vanillin-glucoside, where native degradation genes were removed to prevent carbon loss of APL aromatic metabolites to cell growth. Here, the activity of hcaC/4CL cannot be removed, as the pathway to resveratrol would then be incomplete. Moreover, removing the subsequent step in the pathway, hcaA (Supplemental Figure S19), would trap ferulate in its degradation at 4-hydroxy-3-methoxyphenyl-β-hydroxypropyl-CoA (feruloyl-CoA) and would cause a potentially toxic metabolite buildup.

We identified vanillin synthase of *Vanilla planifolia*^29^ as a potential solution to this issue. Vanillin synthase directly converts ferulate to vanillin, and previous studies have shown it is selective for ferulate, and does not act on *p*-coumarate or another related aromatic metabolite caffeic acid^29^. We incorporated vanillin synthase by plasmid expression into Δ*hcaC*. Though this strain showed a statistically significant decrease in ferulate concentration (p-value 0.0042), and an unexpected statistically significant increase in *p*-coumarate concentration (p-value 0.00245), the activity was weak resulting in only a 4% decrease in ferulate concentration (Figure S20). We hypothesize the increase in *p*-coumarate may be due to promiscuous activity of downstream catabolic enzymes for ferulate degradation that can cleave the O-methyl group at the meta position (*vanAB*) and that may be able to cleave the subsequent meta hydroxy group, thus creating *p*-coumarate from ferulate. Though promising, this strategy will likely only be effective if vanillin synthase can be engineered for superior turnover, and thus was not further pursued in this study.

Having established that ADP1’s native metabolism carries out the first necessary step to convert *p*-coumarate to resveratrol, we added the remaining enzyme, resveratrol synthase (STS) from *Vitis vinifera*. We compared chromosomal and plasmid-based expression in the context of supplying only *p*-coumarate or in full mock APL. These experiments showed resveratrol production in both media contexts, but only with plasmid-based STS expression (Figure 5). Chromosomal expression presumably provided insufficient STS expression. While a greater titer was obtained from *p*-coumarate feeding alone (13.1 mg/L), full APL did show resveratrol production (5.4 mg/L), thus demonstrating ADP1 can successfully synthesize another natural product class (flavonoid/polyketide) from mock APL.

## Conclusion

This work demonstrates the unique potential for *Acinetobacter baylyi* ADP1 within sustainable chemistry, specifically with respect to the biotechnological upgrading of lignin feedstocks. ADP1 combines the metabolic utility of the β-ketoadipate pathway with facile genetic engineering. ADP1’s engineerability is clearly demonstrated in the >20 successive, scarless genetic knockouts, including expression of eight separate heterologous enzymes (COMT, UGT, mtn, luxS, metK, CysE*, metA*, and metB) simultaneously. Of note, all of the necessary cloning to generate these strains was carried out by a single graduate student over the course of one year.

This scale of cloning was enabled by a combination of ADP1’s native homologous recombination, natural competency, and our recently developed markerless and scarless Cas9-based counterselection method^23^. Such a scope of genetic modification would be challenging for even well-established hosts such as *E. coli* and *S. cerevisiae*, and this work provides only a glimpse of the potential for synthetic biology applications. In addition to the facile cloning, ADP1 grows quickly and doesn’t require specialize media or reagents, and thus could be readily adopted by any lab that already conducts *E. coli* cloning. Although these strengths are promising, it is worth mentioning that this work suggests a heterologous expression capacity restraint in ADP1, which is worth further exploration. In addition, it remains to be seen how ADP1, a strict aerobe, will perform at scale. ADP1 is one among several microbial hosts being explored for lignin valorization, (*Pseudomonas putida*^48–50^ and *Rhodoccocus jostii*^51^ representing two others), and each of these microbes have advantages to offer. It remains to be seen if a single host will emerge as ideal. Continued development of each in the near-term will allow for more flexibility as these determinations are made, especially if synthetic microbial communities are to be explored^9^.

The upgrading of aromatic lignin monomers to aromatic products via the metabolic funneling approach is also an important demonstration, specifically our use of ADP1 as a “molecular sieve” where only selected lignin aromatic species are retained. Others have recently suggested taking a more atom economic approach to lignin upgrading^52^. However, previous examples have not used complex feedstock mixtures and included heterologous biosynthetic steps, like shown here. This work clearly demonstrates this potential.

Lastly, the natural products made in this study have value in their own right. Vanilla is the most utilized flavor compound globally, and its current supply is primarily provided by chemical synthesis from petroleum derived compounds^53^. Consumer preferences disfavor this approach, yet the alternative of natural sourcing via *Vanilla plantifolia* is not feasibly scalable. Biotechnological production, as others have pursued^54–56^, provides a possible middle ground for obtaining classifiably natural vanilla via a sustainable and renewable process. Utilizing lignin for this application, which already contains vanillin and other chemically proximal species, offers a synthetically advantaged (fewer biochemical steps) route. Even traditional chemical approaches are looking to leverage vanillin synthesis directly from lignin^57^. Likewise, resveratrol is a desirable nutraceutical, and utilizing the *p*-coumarate in mock APL provides a biochemically advantaged route for its production. Both approaches circumvent the need to take a feedstock down through glycolysis (or another catabolic pathway) and then back up through shikimate or chorismate biosynthesis (and beyond). This study provides a key demonstration for taking greater advantage of lignin’s chemical potential, which will be essential for achieving a biomass-based bioeconomy and will help move towards closed loop carbon cycles via waste valorization.

## Data Availability

Cloning files for all genetic constructs are provided with the supplementary materials as Genebank files.

## Conflict of Interest

B.W.B. and K.E.J.T. have filed a patent (US Patent App. 17/681,434) for vanillin-glucoside synthesis with ADP1 from lignin.

## Acknowledgements

Sanger sequencing was supported by the Northwestern University NUSeq Core Facility. B.W.B. was supported in part by a National Institutes of Health Training Grant (T32GM008449) through Northwestern University’s Biotechnology Training Program). National Science Foundation (NSF) Collaborative Grant [MCB 1614953 to K.E.J.T.].

