## Supplemental Information for "Aromatic natural products synthesis from aromatic lignin monomers using *Acinetobacter baylyi* ADP1"

### Table of Contents

Methods and Materials……………………………………………………………………………………………………..3-5

Supplemental Figures…………………………………………………………………………………………………………6-17

Supplemental Tables………………………………………………………………………………………………………..18-20

References…………………………………………………………………………………………………………………………21

### Materials and Methods

*Cloning*

Cloning methods were carried out as described in extensive detail in a previous work[1]. All ADP1 cloning was conducted with the "ISx" strain of ADP1, generously provided by the lab of Jeffrey Barrick, which has the insertion sequences removed[2]. The enzymes used in this study were obtained as follows, Catechol O-methyltransferase (COMT) from *Homo sapiens* (Uniprot P21964) was codon optimized for *E. coli* and synthesized by IDT. The amino acid sequence for UGT72E2 of *Arabidopsis thaliana* was taken from Uniprot (Q9LVR1), codon optimized for *E. coli*, and synthesized by IDT. Car with the partner Ppt was obtained from previous work in our laboratory[3]. The sequence for Car with the partner Sfp was taken from the literature[4] and built as an operon as described therein. This sequence was codon optimized for ADP1[5] and synthesized as an operon by Twist Biosciences. The amino acid sequence for vanillyl-alcohol oxidase (VAOX) from *Penicillium simplicissimum* was taken from Uniprot (P56216), codon optimized for ADP1, and synthesized by Twist Biosciences. All genes listed were cloned first into pBWB162 (Addgene #140634) in the place of mCherry by Gibson assembly.

Knock out and knock in methods were carried out in the same manner as those described in a previous work[1] concerning the Cas9-based scar-less markerless transformation. However, one adaptation was made, which involved the utilization of a universal gRNA and a landing pad. By creating a gRNA that was validated to cut at the antibiotic resistance gene KanR (sequence GCCGTCTAAGCTATTCGTAT, replacing the pcaHG gRNA in pBWB419 [Addgene #140637]), a landing pad of KanR-lacI-Trc-mCherry was used repeatedly for each successive knock out or knock in, precluding the need to design, generate, and validate a new gRNA for each new locus. Because of this change in procedure, knock out and knock in transformations were no longer carried out in one-step, but involved a two-step process of introducing the landing pad and then later removing it either with nothing (knock out) or a new gene of interest. Cloning files of the different loci with the landing pad knocked in have been provided with the supplemental materials.

For the SAM enzyme integration tests, each *E. coli* enzymes was first amplified by PCR from the *Escherichia coli* (K12 MG1655) chromosome and incorporated into pBAV1k-KanR-lacI-Trc by Gibson assembly. This plasmid was used as the template for overlap PCR for integrations into the ADP1 genome, for which each enzyme bore the Trc promoter and the agga RBS. For the six enzymes: *luxS* was integrated at the *calB* locus, *mtn* was integrated at *quiA*, *metK* was integrated at *ACIAD2929*, *metB* was integrated at *ACIAD1577-1578*, the feedback resistant version of *metA* (*metA**) was integrated at *acoD*, and the feedback resistant version of *CysE* (*CysE**) was integrated at *entA* (also known as *dbhA*). Feedback resistant mutants were created by PCR mutagenesis using the NEB KLD kit. CysE mutagenesis was carried out in one step, metA mutagenesis was carried out in two steps.

For resveratrol cloning, the enzymes 4CL and STS were obtained from Addgene (#139790)[6]. These enzymes were cloned into pBAV1k-kanR-lacI-Trc by Gibson assembly without further modification. The sequence for Vanillin Synthase of *Vanilla planifolia* was obtained from Uniprot (A0A0F7G352). The amino acid sequence was codon optimized for ADP1[5] and the optimized sequence was synthesized by Twist Biosciences and cloned into pBAV1k-kanR-lacI-Trc by Gibson assembly. Knock out and knock in procedures were carried as described above. STS was integrated as the cassette “lacI-Trc-STS”, with the “agga” RBS at *ACIAD1577-1578*. Plasmid maps and integration files can be found with the supplemental materials, along with an Excel file with all primers used in the study.

*Culturing*

General culturing and induction methods were conducted as described previously[1]. For the vanillin enzyme test and vanillin-glucoside production cultivations, overnight cultures inoculated from glycerol stocks were grown in LB at 30°C, with the appropriate antibiotic. The next morning, these pre-cultures were sub-cultured 1:100 into 3 mL of Difco M9 minimal medium with trace metal added. Trace metal solution was prepared as described in a prior work[7]. Cultures were typically induced from 0 hrs with 1 mM IPTG. Substrate provisions for enzymes tests were typically given a 1 mM. However, as vanillyl alcohol was degraded quickly, it was provided at 2 mM for the vanillyl alcohol oxidase (VAOX) test. In addition, though toxic, PCA was provided at 1.5 mM in subsequent tests of COMT activity, as previous tests had shown exhaustion of the 1 mM pool. For the mock APL medium, Difco M9 minimal medium was used in a 24-hr cultivation, with trace elements, and carbon was provided as follows: 1% glucose, 1 g/L acetate, 1.5 mM p-coumarate (250 mg/L), 0.5 mM ferulate (97 mg/L), 0.5 mM vanillate (84 mg/L), 0.5 mM p-hydroxybenzoate (69 mg/L), 0.5 mM p-hydroxybenzaldehyde (61 mg/L). The VAOX integration strain cultivation was run for 48-hours.

For resveratrol culturing, pre-cultures for *p*-coumarate and ferulate degradation and resveratrol production were inoculated from a glycerol stock and grow in LB overnight at 30°C, 250 rpm. The next morning, cultivations were diluted 1:100 into Difco M9 minimal medium with trace metals[7]. For the degradation test, 1 mM p-coumarate (163 mg/L) and 1 mM ferulate (194 mg/L) were supplied, along with 1% glucose and 1 g/L acetate. Cultures were grown overnight at 30°C, 250 rpm. Antibiotics (kanR) were used for plasmid containing cultures and IPTG was supplied where necessary for induction at 1 mM concentration. For resveratrol production, a similar approach was used, but for production from *p*-coumarate 4 mM (652 mg/L) *p*-coumarate was supplied, along with 1% glucose and 1 g/L acetate. For APL conditions the M9 minimal medium with trace metals contained, 5 g/L glucose, 1 g/L acetate, 1.5 mM p-coumarate (250 mg/L), 0.5 mM ferulate (97 mg/L), 0.5 mM vanillate (84 mg/L), 0.5 mM p-hydroxybenzoate (69 mg/L), 0.5 mM p-hydroxybenzaldehyde (61 mg/L), and 0.5 mM syringate (106 mg/L). Antibiotics were supplied for plasmid containing cultures and induction was carried out from 0 hr with 1 mM IPTG.

*Vanillin HPLC*

After overnight incubation, and $OD_{600}$ values were taken, cultivations were centrifuged at 4°C for 10 min at 4,000 x *g*. The supernatant was filtered with a 0.2 µM filter, and 1 mL was added to glass screw top vials that were sealed with PTFE/silicone caps (Filtrous). Samples were analyzed using an Agilent 1200 Series HPLC equipped with a BioRad HPX-87H chromatography column, an Agilent G1315C Diode Array Detector (DAD), and an Agilent G1362A Refractive Index Detector (RID). The mobile phase was 10% (v/v) acetonitrile, 90% (v/v) 5 mM sulfuric acid (Thermo), pH of approximately 2.3, and an isochromatic method was utilized. The column was equilibrated for 1 hour at a flow rate of 0.600 mL/min and a column temperature of 60°C.

Following equilibration at 0.600 mL/min at 60°C, the following method was used to run samples. The injection volume was 5.00 µL. Run time was 60 minutes with post-time of 1 minute. The autosampler was maintained at a temperature of 4°C and the RID at a temperature of 35°C. The DAD signal at 206 nm was used to quantify all aromatic species, including vanillin-glucoside. Standards obtained from Sigma-Aldrich were run for each aromatic acid to determine retention times and to generate a standard curve for any metabolite where quantification of titer is given.

*Resveratrol HPLC*

After cultivations were run overnight, they were centrifuged for 10 min at 4,000 x *g*. The supernatant was then filtered by with a 0.2 µM filter, and 1 mL was added to glass screw top vials that were sealed with PTFE/silicone caps (Filtrous). Samples were analyzed using an Agilent 1200 Series HPLC equipped with an Agilent ZORBAX Extend-C18 Rapid Resolution HT 4.6 x 100mm, 1.8 micron, an Agilent G1315C Diode Array Detector (DAD), and an Agilent G1362A Refractive Index Detector (RID). The mobile phase was 25% (v/v) acetonitrile, 0.1% formic acid, with a pH of approximately 2.6, and an isochromatic method was utilized. The column was equilibrated for 1 hour at a flow rate of 0.600 mL/min and a column temperature of 30°C. Following equilibration at 0.500 mL/min at 30°C, the following method was used to run samples. The injection volume was 5.00 µL. Run time was 30 minutes with post-time of 1 minute. The autosampler was maintained at a temperature of 4°C and the RID at a temperature of 35°C. The DAD signal at 302 nm was used to quantify p-coumarate, ferulate, and resveratrol. Standards obtained from Sigma-Aldrich were run for each compound to determine retention times and to generate a standard curve for resveratrol.

### Supplemental Figures

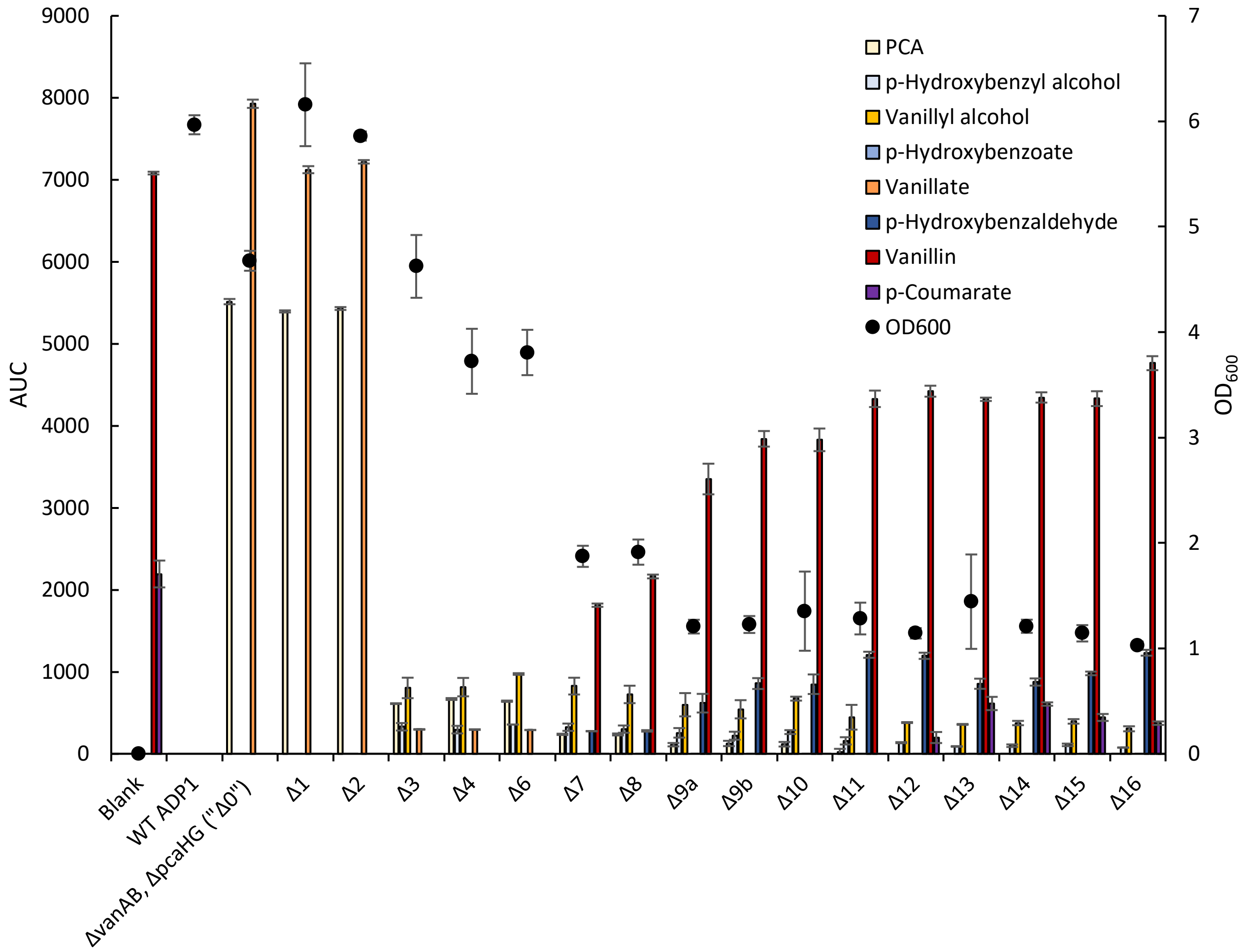


**Figure S1**. *Metabolite retention profiles of the putative vanillin dehydrogenase (VDH) knock out strains.* Δ and number indicate the number of putative vanillin dehydrogenases removed from the strain. Colored bars indicate relative amount (left y-axis) of each metabolite present, error bars are for standard deviation of biological triplicate. Black dots indicate $OD_{600}$ (right y-axis) of each strain, error bars are for standard deviation of biological triplicate.

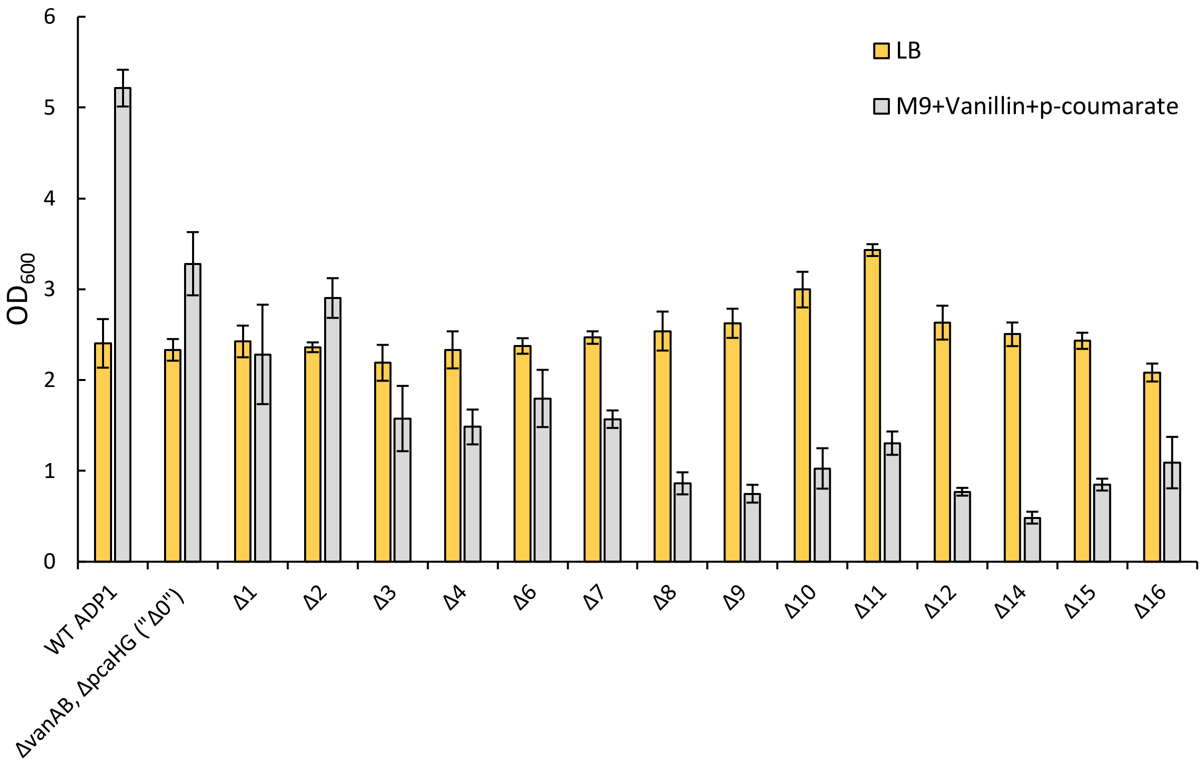


**Figure S2**. *Growth of vanillin dehydrogenase knock out strains on LB and M9.* Yellow bars indicate strain growth ($OD_{600}$) on LB, error bars are standard deviation for biological triplicate. Gray bars are for growth ($OD_{600}$) on M9 with vanillin and *p*-coumarate, error bars are for standard deviation of biological triplicate.

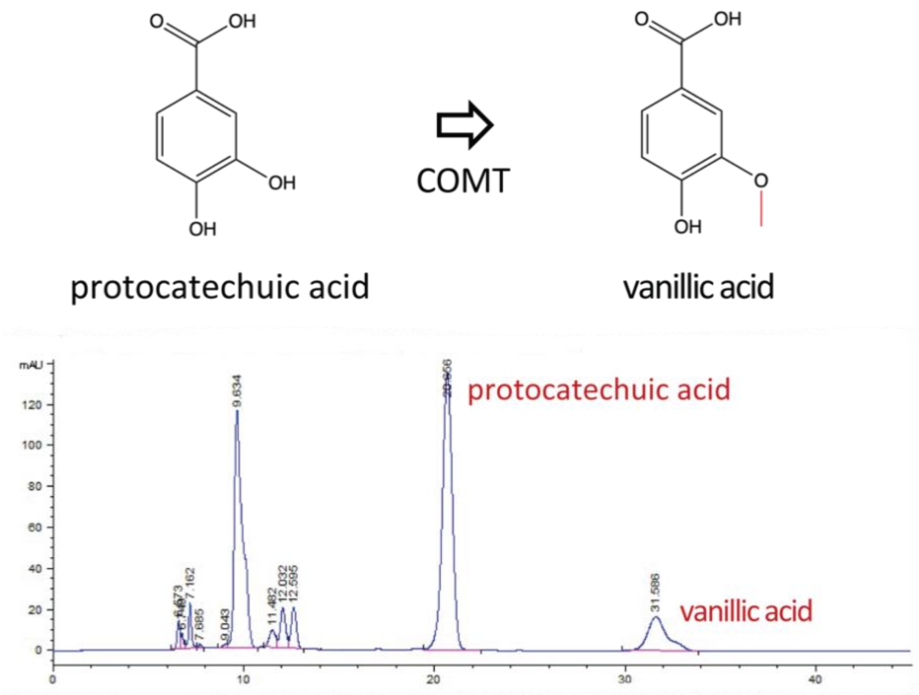


**Figure S3**. *COMT enzymatic activity demonstration.* HPLC chromatogram for the activity test of catechol O-methyltransferase (COMT) of *Homo sapiens* expressed heterologously in ADP1. For this experiment, protocatechuic acid (PCA) was fed to ADP1 containing COMT. As can be seen, vanillic acid (retention 31.586 min) is produced from this cultivation. Retention times determined by standards.

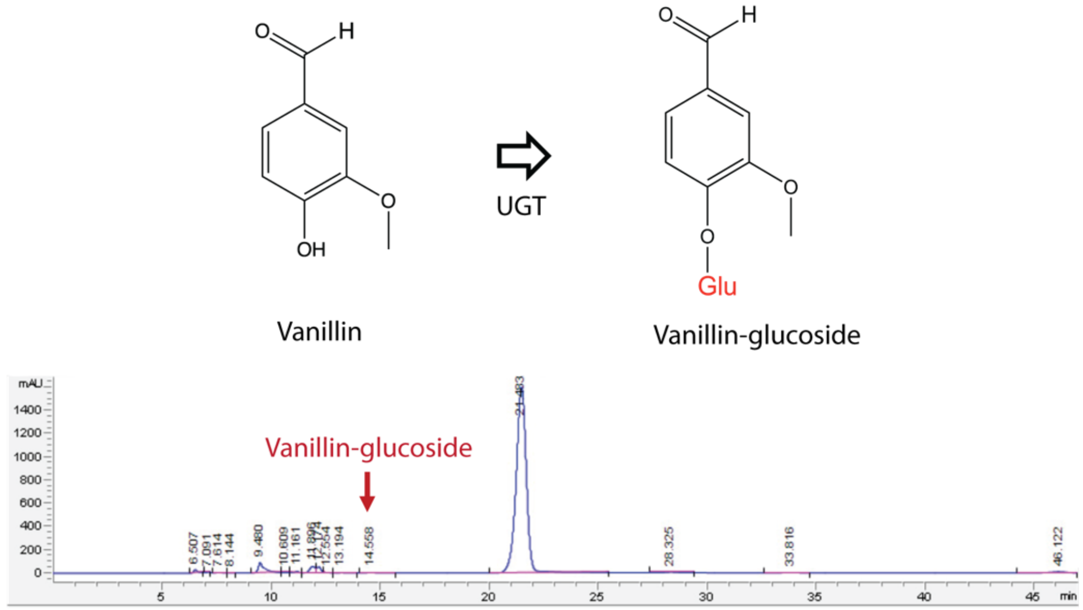


**Figure S4**. *UGT72E2 enzymatic activity demonstration*. HPLC chromatogram depicting the activity of *Arabidopsis thaliana* UDP-glucose dependent glycosyltransferase UGT72E2 as expressed heterologously in ADP1 with vanillin provided. As vanillin is degraded by ADP1, conversion must occur quickly to be observed and only trace vanillin-glucoside is observed (retention time 14.558 min). Retention time determined by standards.

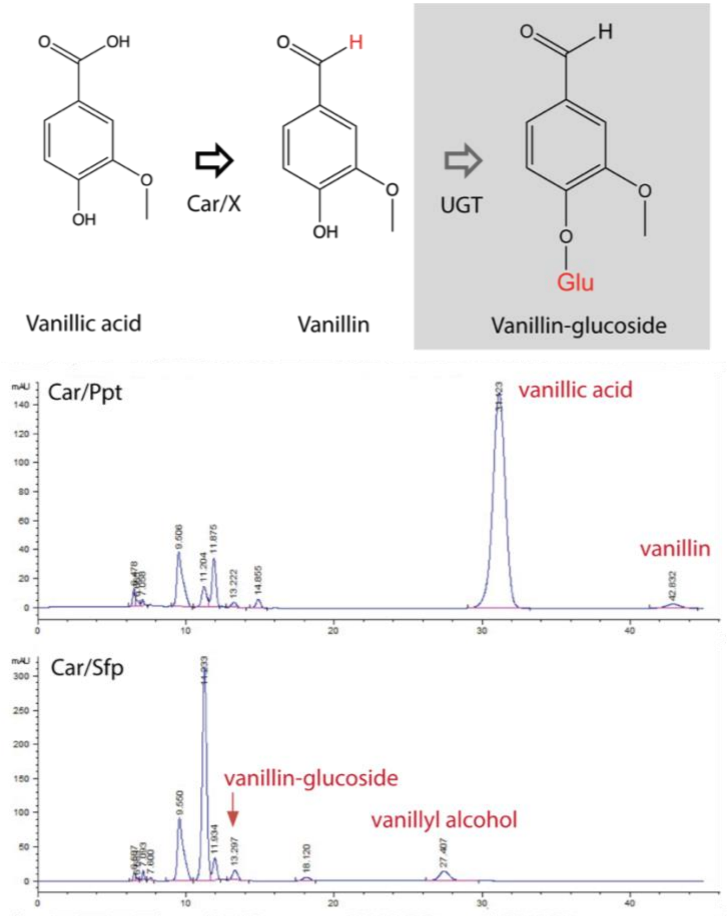


**Figure S5**. *Car enzymatic activity demonstration comparing two different cofactor enzymes.* HPLC chromatogram demonstrating heterologous activity of Car (*Nocardia iowensis*) in ADP1, with vanillate provided and in the context of a strain that also contains UGT72E2. The UGT enzyme was included to help "capture" vanillin conversion in the form of vanillin-glucoside, as vanillin is degraded by ADP1. Two separate partner enzymes were tested to determine whether either provided turnover benefit. While Car/Ppt (upper panel) does show turnover, Car/Sfp exhausted the entire vanillate provided, showing two products of vanillin conversion, vanillyl alcohol (degradation product) and vanillin-glucoside (from UGT conversion).

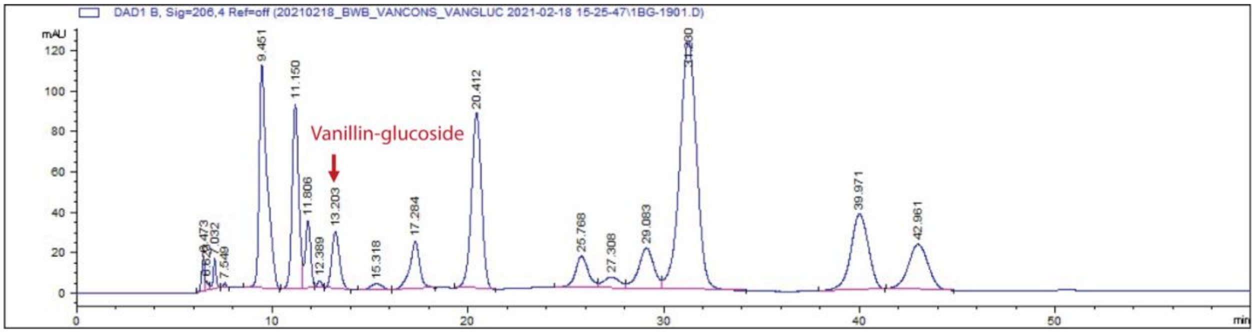


**Figure S6**. *Vanillin-glucoside production from simplified mock APL HPLC chromatogram.* HPLC chromatogram showing results for the initial test for vanillin-glucoside production using "Δ20" as a strain background, where COMT is expressed from the integration of a lacI-Trc-COMT cassette at *vanAB,* UGT72E2 is expressed from a cassette of Trc-BCD9-UGT72E2 integrated at *pcaHG*, and Car/Sfp is plasmid-expressed with pBAV1k-kanR-lacI-Trc-Car/Sfp. The mock APL in this experiment did not contain *p*-hydroxybenzaldehyde or syringate. The cultivation was run in M9, overnight, at 30ºC in M9 minimal medium with trace metals, in a 3 mL culture.

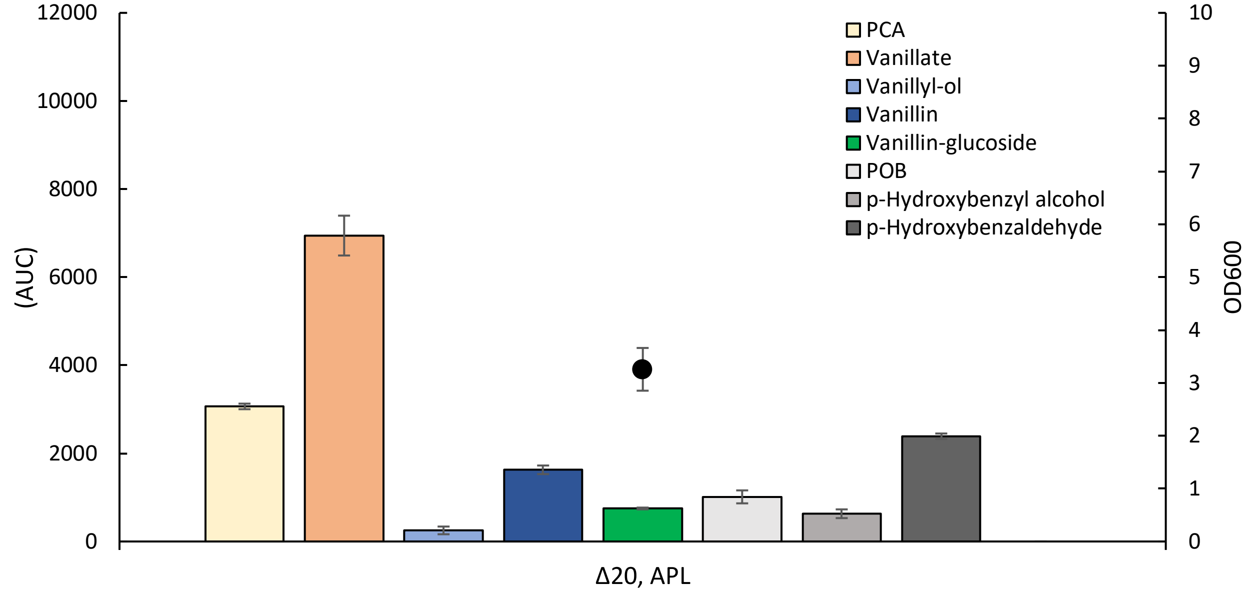


**Figure S7**. *Vanillin-glucoside production from simplified mock APL histogram.* Histogram shows triplicate data of cultivation described in Figure S6 (HPLC trace). Error bars represent standard deviation for biological triplicate. The greatest accumulation of any metabolite is seen at vanillate (orange bar).

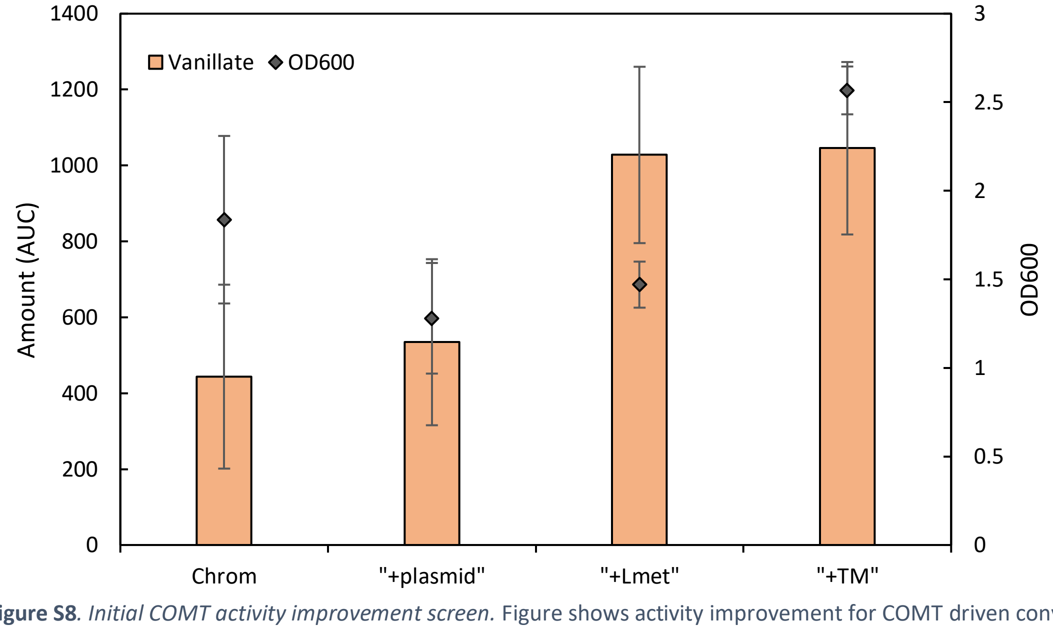


**Figure S8**. *Initial COMT activity improvement screen.* Figure shows activity improvement for COMT driven conversion of PCA to vanillate. Orange bars are amount of vanillate (AUC). Black diamonds represent the optical density ($OD_{600}$). Each cultivation is carried out in the context of the chromosomal integration (*vanAB*::lacI-Trc-COMT) and in M9 minimal medium. “Chrom” is the reference condition with any additions. “+plasmid” represents the addition of plasmid expression from pBAV1k-lacI-Trc-COMT. “+plasmid” and shows a modest improvement in PCA to vanillate conversion, with a concurrent decrease in cell growth ($OD_{600}$). “+Lmet” represents the addition of 10 mM L-methionine to the medium. “+Lmet” shows nearly a doubling of vanillate conversion. The same result was obtained for “+TM”, which represents the addition of the trace metal solution to the base condition. All conditions were provided 1% glucose, 25 mM acetate, and 2 mM PCA. Error bars represent standard deviation of biological triplicate.

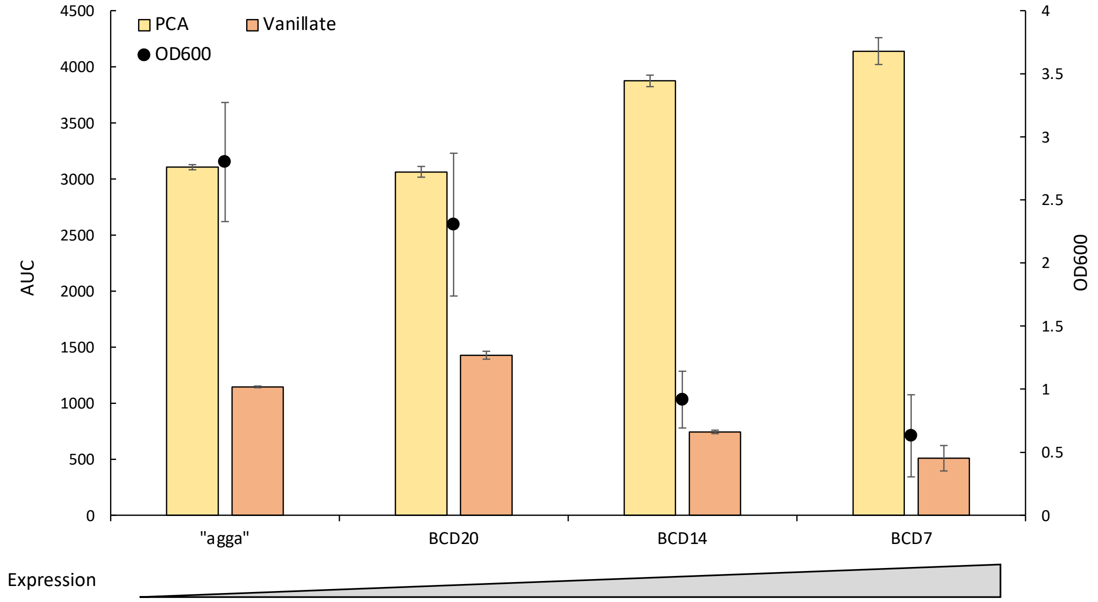


**Figure S9.** *COMT activity improvement from modulating chromosomal expression through ribosomal binding site variants.* Figure shows the change conversion of PCA to vanillate by COMT in different expression contexts. All cases involve chromosomal expression of COMT as a cassette at *vanAB*::lacI-Trc-X-COMT, where X is the ribosomal binding site. To the far left is the reference "agga" ribosomal binding site. As can be seen, expression stronger than BCD20 (BCD14, BCD7) severely impacted cell growth ($OD_{600}$) and negatively impacted conversion to vanillate. However, BCD20 does show improvement in conversion (24%, p-value 0.000167) by finding an optimal expression. Bars are amount of PCA (yellow) and vanillate (orange) respectively (AUC, HPLC trace), and the black circles represent optical density ($OD_{600}$). Error bars are for standard deviation of biological triplicate.

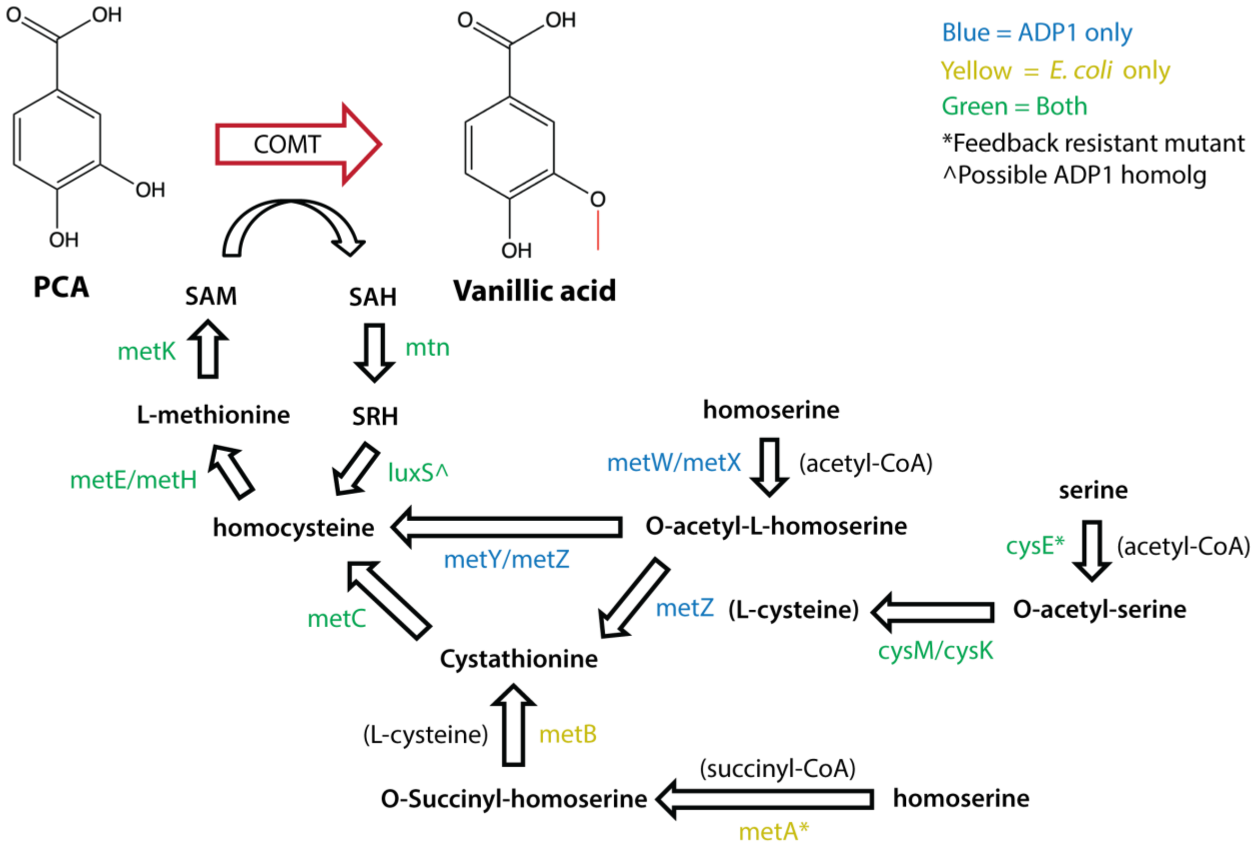


**Figure S10**. *SAM enzyme mapping*. Figure shows routes to s-adenosylmethionine (SAM), the cofactor used by COMT to donate a methyl group to methylate PCA to generate vanillate. In blue are the enzymes (routes) unique to ADP1. In yellow are the enzymes (routes) unique to *E. coli*. In green are the shared steps.

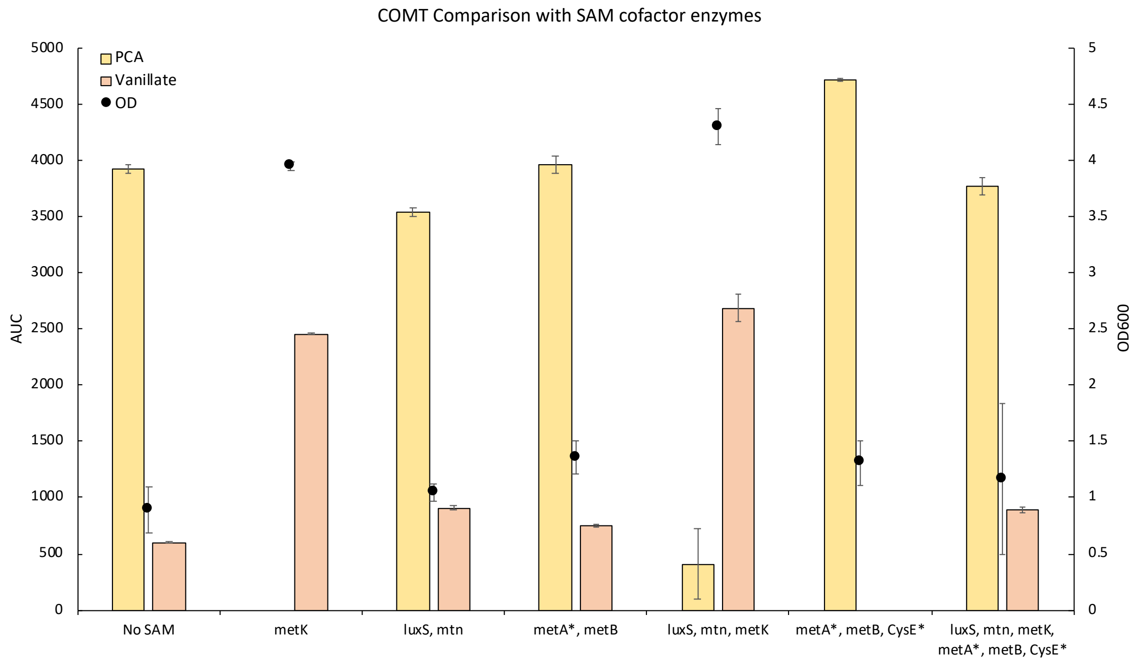


**Figure S11**. *COMT activity improvement from the inclusion of SAM cofactor pool enzymes.* Figure shows the improvement of COMT activity in converting PCA to vanillate with the inclusion of different grouping of SAM pool enzymes (Figure S10 depicts their roles). On the far left is the reference strain with no modifications to the SAM pool enzymes. The inclusion of *metK* alone ("metK) shows a strong improvement (4.1-fold) in COMT conversion and greatly improves cell growth ($OD_{600}$). While other combinations provide some benefit, the only combination with greater activity improvement than *metK* alone is that with *metK*, *mtn*, and *luxS* (4.48-fold improvement). A strain with all six tested enzymes added showed lower COMT activity than either *metK* alone or *metK*, *mtn*, and *luxS*. Error bars represents standard deviation from biological triplicate.

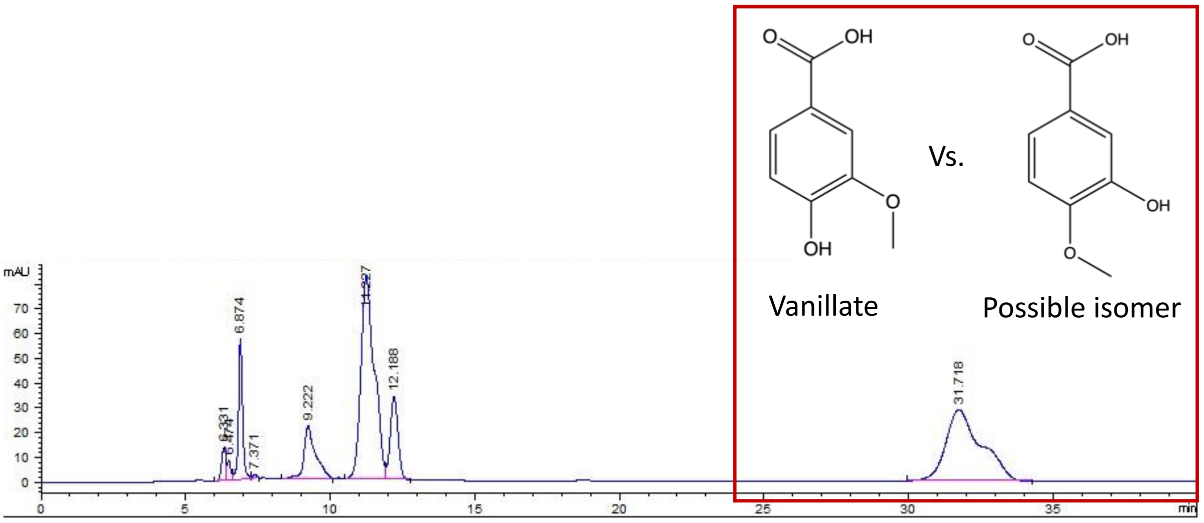


**Figure S12**. *Improved COMT activity HPLC chromatogram.* Improved COMT activity from the inclusion of the SAM pool enzymes (*luxS*, *mtn*, and *metK*) shows much greater conversion of PCA to vanillate, however the peak eluting at the time of vanillate now contains a shoulder that may represent a vanillate isomer "isovanillin", potentially generated by the promiscuous methylation of the *p*-hydroxy group of PCA.

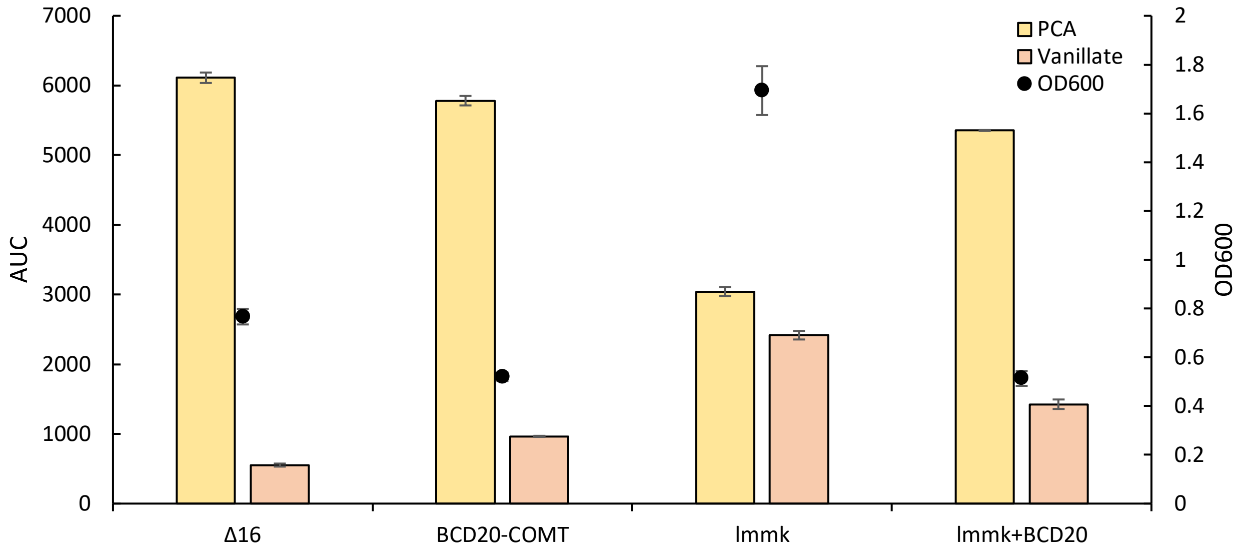


**Figure S13***. Testing COMT improvements and their combination.* Figure shows the improvements in COMT turnover of PCA to vanillate in different strain backgrounds. First is "Δ16" as a reference, a strain with COMT integrated as *vanAB*::lacI-Trc-COMT. Next is "BCD20-COMT", which is a modification from "Δ16" only with respect to the RBS used for COMT and was the best variant from the prior RBS-based expression modification testing. Third is "lmmk", which represents the best combination of SAM enzymes from the prior screen (*luxS*, *mtn*, and *metK*). Last is "lmmK+BCD20", which represents the combination of the two prior optimum strains. Error bars represent standard deviation from biological triplicate.

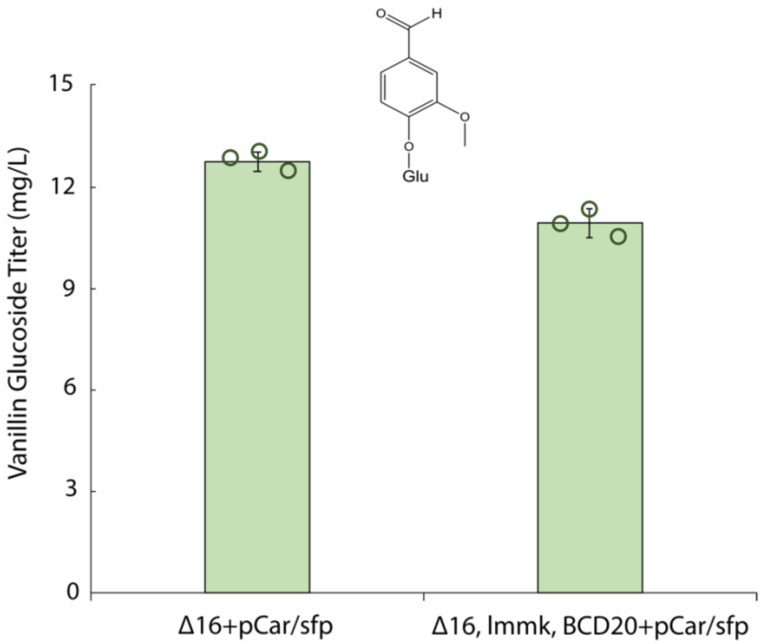


**Figure S14.** *Vanillin-glucoside production.* Figure shows the comparison between the base Δ16 strain with Car/Sfp provided by plasmid expression and the strain with the COMT optimizations (expression and SAM cofactor supply).

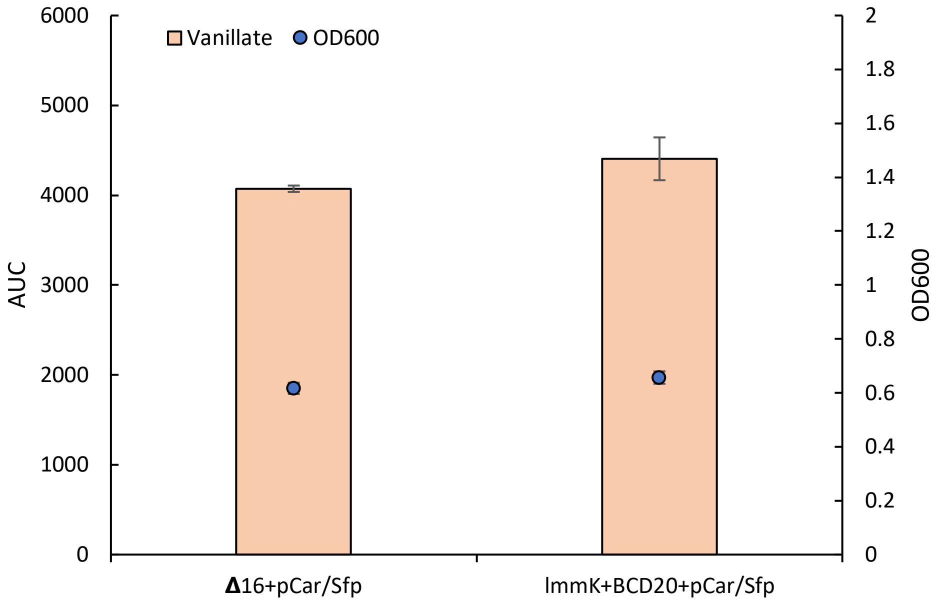


**Figure 15.** *Vanillate difference in COMT optimization strain in the context of APL and full vanillin-glucoside pathway production.* The strain with the COMT optimization does show a modest (8%) increase in vanillate compared to the reference strain. P-value for a student's t-test, two tailed, equal variance is 0.074. Error bars represent standard deviation of biological triplicate.

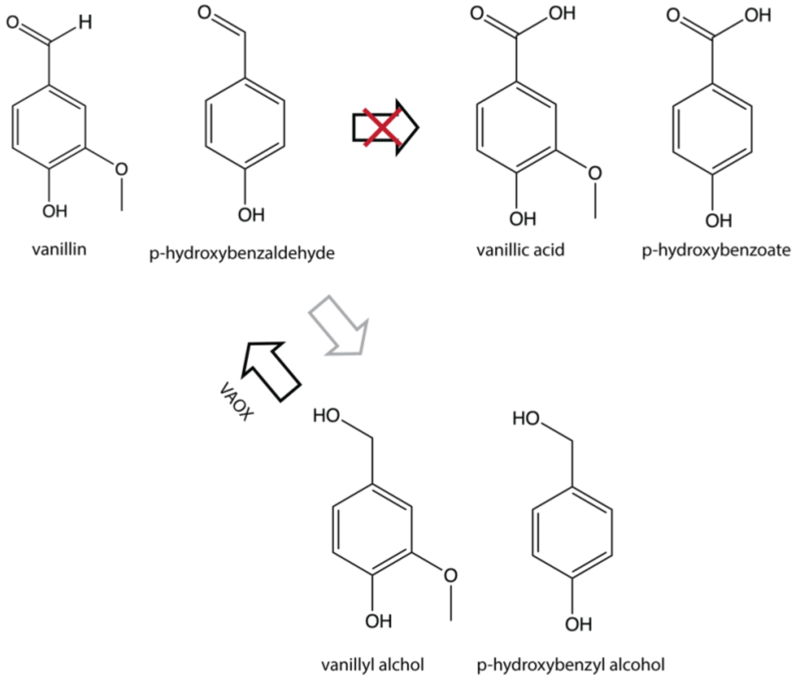


**Figure S16***. Expected activity of putative vanillin dehydrogenases on vanillin and p-hydroxybenzaldehyde.* As known for *p*-coumarate and ferulate's degradation in *P. putida* (MetaCyc), where a single enzyme (*vdh*) is shared for the degradation of vanillin and *p*-hydroxybenzaldehyde, it is proposed that the same promiscuous enzymes in ADP1 capable of degrading vanillin are likely also able to degrade *p*-hydroxybenzaldehyde. In addition, from the literature it is know that the same set of enzymes in *E. coli* had dehydrogenase activity on both vanillin and benzaldehyde [8].

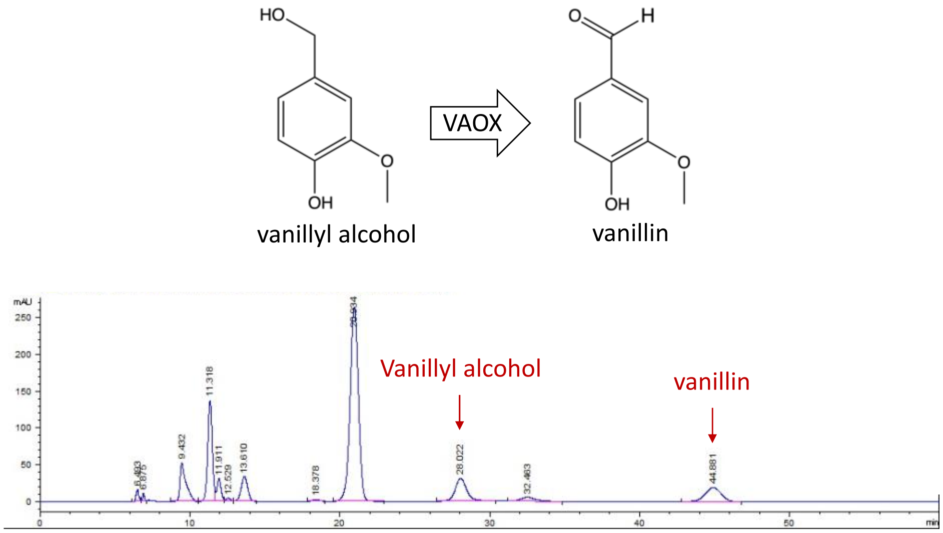


**Figure S17**. *Vanillyl alcohol oxidase demonstration.* HPLC chromatogram shows conversion of vanillyl alcohol to vanillin by vanillyl alcohol oxidase (VAOX) of *Penicillium simplicissimum* in the context of the "Δ16" ADP1 strain fed vanillyl alcohol and *p*-hydroxybenzoate. *p*-hydroxybenzoate feeding was used to help slow vanillyl alcohol degradation by kinetically competing for degradation enzymes long enough to observe VAOX activity.

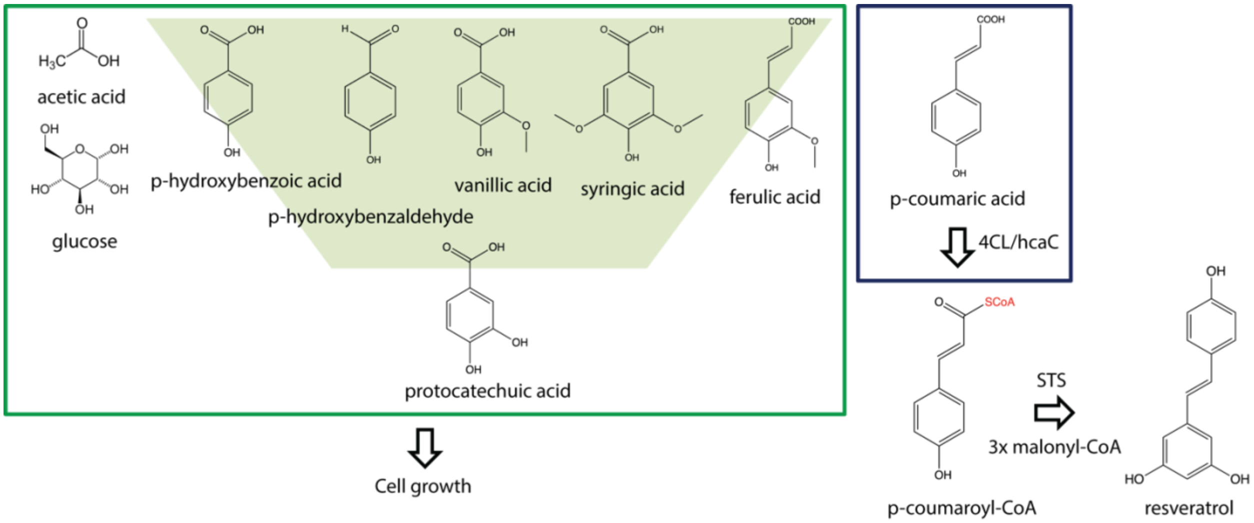


**Figure S18.** *Resveratrol production from mock APL.* Figure depicts strategy for utilizing ADP1 to convert mock APL to resveratrol. In this approach, *p*-coumarate is converted to *p*-coumaroyl-CoA by a CoA ligase (4CL or hcaC) and then converted by resveratrol synthase (STS) to resveratrol using 3 units of malonyl-CoA, while the rest of the carbon in APL is converted to cell growth via central carbon metabolism, including the utilization of the β-ketoadipate pathway.

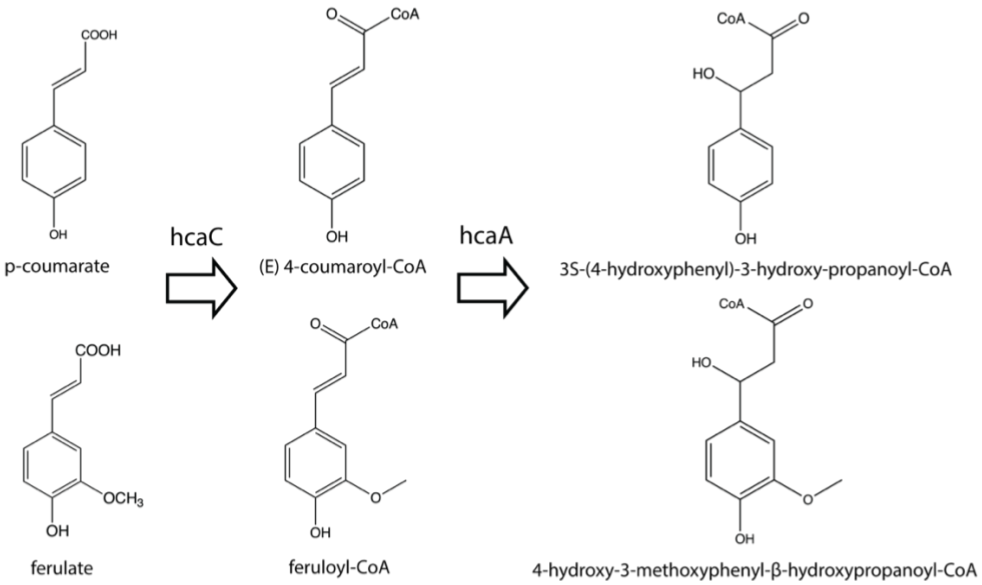


**Figure S19.** *ADP1 native p-coumarate and ferulate metabolism first steps.* The degradation pathways for *p*-coumarate and ferulate share their first two steps in ADP1 with *hcaC* performing the initial CoA ligation and *hcaA* performing the subsequent hydroxylation.

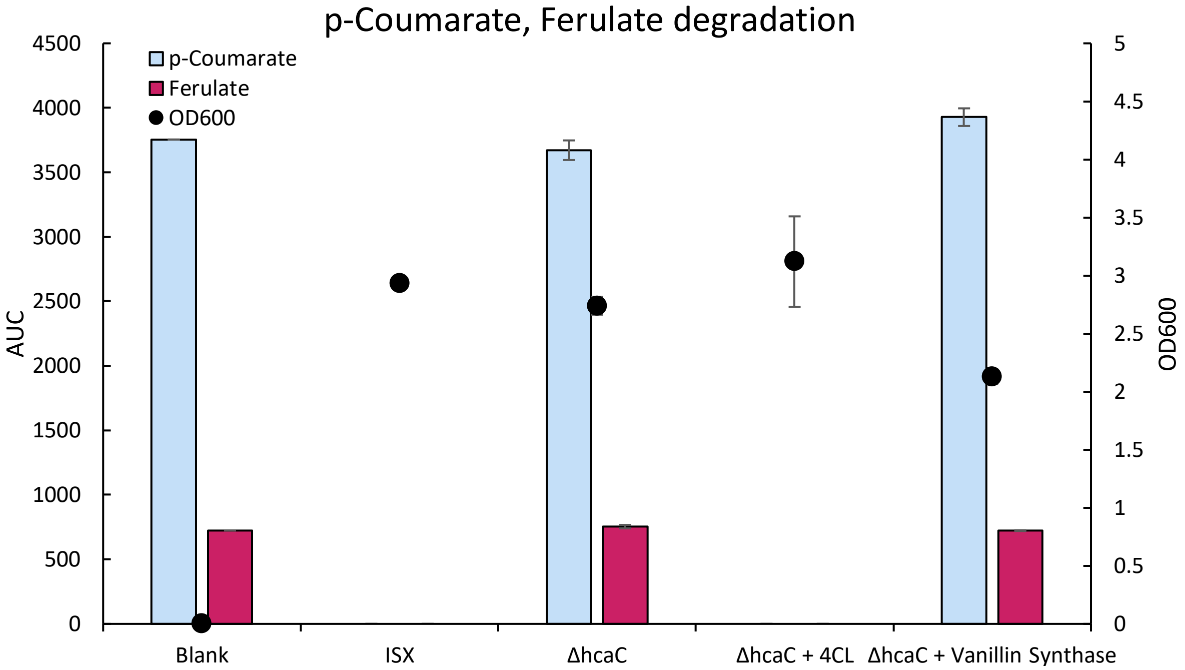


**Figure S20**. *ADP1 degradation of p-coumarate and ferulate*. Figures shows the degradation of 1 mM *p*-coumarate and 1 mM ferulate supplied simultaneous by various ADP1 strains. The first condition is the blank medium measurement by HPLC, given as a reference. Next is "ISx", a wild-type ADP1 strain with its insertion sequences removed[2], which shows complete consumption of *p*-coumarate and ferulate. By deleting *hcaC* ("ΔhcaC"), ADP1 now retains both *p*-coumarate and ferulate. By taking "ΔhcaC" and supplementing it with 4-coumarate-CoA ligase (4CL), ADP1 is once again able to metabolize *p*-coumarate and ferulate. By supplying "ΔhcaC" with vanillin synthase, a slight (4%), but statistically significant (p-value of 0.0042 for student t-test, two tailed, and equal variance) decrease is observed for ferulate, along with a statistically significant (p-value of 0.00245) increase in *p*-coumarate. Error bars are standard deviation for biological triplicate, expect for vanillin synthase which contained one additional replicate (4 total).

### Supplemental Tables

**Table S1**. *List of putative vanillin dehydrogenases in ADP1*. List determined by psi-BLAST searches on known vanillin dehydrogenase from *Pseudomonas putida*, *Corynebacterium glutamicum*, and the promiscuous dehydrogenases found in a prior study[8]. "Disp." indicates whether a gene was able to be removed in a prior ADP1 single-gene knockout library study[9]. "Succ." column indicates relative expression from growth on succinate. "Quin." indicates relative expression from growth on quinoate. "Fold" indicates the fold increase in expression from growth on quinoate compared to succinate, with quinoate acting as a proxy for aromatic metabolite induction of gene expression, as determined by a previous study[10]. "Hom" indicates homology % to the gene that was submitted for the psi-BLAST search. The final column indicates which gene was used as a seed in the psi-BLAST.

| Name | ACIAD | Activity | Disp. | Succ. | Quin. | Fold | Hom | Source |
|---|---|---|---|---|---|---|---|---|
| *hcaB* | ACIAD1725 | hydroxybenzaldehyde dehydrogenase | Yes | 118 | 267 | 2.263 | 62.32 | *P. putida vdh* |
| *areC* | ACIAD1430 | benzaldehyde dehydrogenase II | Yes | 41 | 575 | 14.024 | 37.12 | *P. putida vdh* |
| *areB* | ACIAD1429 | aryl-alcohol dehydrogenase | Yes | 42 | 738 | 17.571 | 26.84 | *E. coli yahK* |
| *calB* | ACIAD0503 | coniferyl aldehyde dehydrogenase (CALDH) | Yes | 4135 | 3045 | 0.736 | 30.7 | *P. putida vdh* |
| | ACIAD1577 | putative aldehyde dehydrogenase | Yes | 51 | 257 | 5.039 | 31.92 | *P. putida vdh* |
| | ACIAD1578 | putative aryl-alcohol dehydrogenase (Benzyl alcohol dehdyrogenase) | Yes | 37 | 119 | 3.216 | 26.13 | E. coli yahK |
| *betB* | ACIAD1009 | NAD+-dependent betaine aldehyde dehydrogenase | Yes | 405 | 498 | 1.23 | 38.8 | *C. glutamicum* Ald |
| *quiA* | ACIAD1716 | quinate/shikimate dehydrogenase | Yes | 2295 | 3.99E+04 | 17.386 | - | |
| *acoD* | ACIAD2018 | aldehyde dehydrogenase. Acetaldehdye dehydrogenase II | Yes | 2885 | 4417 | 1.531 | 68.45 | *C. glutamicum* Ald |
| *frmA* | ACIAD1879 | Alcohol dehydrogenase class 3 | Yes | 1577 | 2860 | 1.814 | 26.63 | *E. coli yahK* |
| *adhA* | ACIAD3339 | alcohol dehydrogenase, cinnamyl alcohol dehydrogenases | Yes | 3280 | 953 | 0.291 | 32.76 | *E. coli yahK* |
| dhbA | ACIAD2774 | 2,3-dihydro-2,3-dihydroxybenzoate dehydrogenase "entA" | | 21 | 13 | 0.619 | | |
| | ACIAD3612 | NADP-dependent alcohol dehydrogenase, cinnamyl alcohol dehydrogenase | Yes | 193 | 356 | 1.845 | 58.11 | *E. coli yjgB* (*ahr*) |
| | ACIAD2015 | putative alcohol dehydrogenase | Yes | 375 | 233 | 0.621 | 27.37 | *E. coli yqhD* |
| | ACIAD2929 | putative alcohol dehydrogenase | Yes | 550 | 2299 | 4.18 | 23.88 | *E. coli_yqhD* |
| | ACIAD1743 | putative oxydoreductase protein | Yes | 590 | 1325 | 2.246 | 23.02 | *E. coli_yahK* |
| *tgnC* | ACIAD2542 | Putative aldehyde dehydrogenase | Yes | 240 | 489 | 2.038 | 37.27 | *C. glutamicum* Ald |
| *alrA* | ACIAD3616 | aldehyde reductase | Yes | 244 | 213 | 0.873 | 62.55 | *E. coli dkgB* |
| | ACIAD1261 | putative oxidoreductase, aldo/keto reductase family | Yes | 212 | 491 | 2.316 | 27.06 | E. *coli_yeaE* |
| | ACIAD1950 | putative iron-containing alcohol dehydrogenase | Yes | 961 | 658 | 0.685 | 26.38 | *E. coli yqhD* |
| | ACIAD3642 | Putative aldehyde dehydrogenase | No | 549 | 496 | 0.903 | 28.75 | *P. putida vdh* |
| | ACIAD0998 | Putative aldehyde dehydrogenase. Energy metabolism | Yes | 19 | 27 | 1.421 | 33.81 | *C. glutamicum* Ald |
| | ACIAD3128 | putative oxidoreductase | Yes | 18 | 27 | 1.5 | 26.12 | *E. coli dkgB* |
| | ACIAD1596 | putative oxidoreductase | Yes | 42 | 57 | 1.357 | 25.56 | *E. coli dkgB* |
| *dkg* | ACIAD3281 | 2,5-diketo-D-gluconate reductase | Yes | 2.92E+04 | 1.96E+04 | 0.671 | 39.37 | *E. coli_dkgA* |
| | ACIAD0546 | putative NADP-dependent glyceraldehyde-3-phosphate dehydrogenase | Yes | 2.02E+05 | 2.65E+05 | 1.312 | 29.31 | *P. putida vdh* |

| | | | | | | | | |
|---|---|---|---|---|---|---|---|---|
| | ACIAD1021 | putative (R,R)-butanediol dehydrogenase, acetoin reductase | Yes | 194 | 564 | 2.907 | 26.56 | *E. coli yahK* |
| putA | ACIAD1646 | delta-1-pyrroline-5-carboxylate dehydrogenase | | | | | 27.52 | *P. putida vdh* |
| | ACIAD0131 | 2-ketoglutarate semialdehyde dehydrogenase | | | | | 25.82 | *P. putida vdh* |
| *benD* | ACIAD1437 | cis-1,2-dihydroxycyclohexa-3,5-diene-1-carboxylate dehydrogenase | Yes | 30 | 25 | 0.833 | - | |
| *gabD* | ACIAD2539 | Succinate-semialdehyde dehydrogenase | Yes | 444 | 410 | 0.923 | 32.56 | *P. putida vdh* |
| *gabD* | ACIAD3445 | NADP+ dependent succinate semialdehyde dehydrogenase | Yes | 52 | 52 | 1 | 35.66 | *P. putida vdh* |
| | ACIAD0960 | Succinate semialdehyde dehydrogenase [NADP+] (GabD-like) | Yes | 190 | 188 | 0.989 | 31.14 | *P. putida vdh* |
| *astD* | ACIAD1287 | succinylglutamic semialdehyde dehydrogenase | Yes | 102 | 210 | 2.059 | 27.56 | *P. putida vdh* |
| *mmsA* | ACIAD1604 | methylmalonate-semialdehyde dehydrogenase, oxidoreductase protein | Yes | 131 | 286 | 2.183 | 28.3 | *P. putida vdh* |

**Table S2**. *List of ADP1 knock outs tested for vanillin-glucoside production.* Green indicates genes that were knocked out in the final strain. Blue indicates putative vanillin dehydrogenases that were tested but found to not make a difference for vanillin degradation.

| Name | ACIAD | Activity | Reason |
|---|---|---|---|
| pcaH | ACIAD1711 | protocatechuate 3,4-dioxygenase beta chain (3,4-PCD) | protocatechuate degradation |
| pcaG | ACIAD1712 | protocatechuate 3,4-dioxygenase alpha chain (3,4-PCD) | protocatechuate degradation |
| vanB | ACIAD0979 | vanillate O-demethylase oxidoreductase (Vanillate degradation ferredoxin-like protein) | vanillate degradation |
| vanA | ACIAD0980 | vanillate O-demethylase oxygenase subunit (4-hydroxy-3-methoxybenzoate demethylase) | vanillate degradation |
| *hcaB* | ACIAD1725 | hydroxybenzaldehyde dehydrogenase | vanillin degradation |
| *areC* | ACIAD1430 | benzaldehyde dehydrogenase II | vanillin degradation |
| *areB* | ACIAD1429 | aryl-alcohol dehydrogenase | vanillin degradation |
| *calB* | ACIAD0503 | coniferyl aldehyde dehydrogenase (CALDH) | vanillin degradation |
| | ACIAD1577 | putative aldehyde dehydrogenase | vanillin degradation |
| | ACIAD1578 | putative aryl-alcohol dehydrogenase (Benzyl alcohol dehdyrogenase) | vanillin degradation |
| *betB* | ACIAD1009 | NAD+-dependent betaine aldehyde dehydrogenase | vanillin degradation |
| *quiA* | ACIAD1716 | quinate/shikimate dehydrogenase | vanillin degradation |
| *acoD* | ACIAD2018 | aldehyde dehydrogenase. Acetaldehdye dehydrogenase II | vanillin degradation |
| *frmA* | ACIAD1879 | Alcohol dehydrogenase class 3 | vanillin degradation |
| *adhA* | ACIAD3339 | alcohol dehydrogenase, cinnamyl alcohol dehydrogenases | vanillin degradation |
| dhbA | ACIAD2774 | 2,3-dihydro-2,3-dihydroxybenzoate dehydrogenase (entA) | vanillin degradation |
| | ACIAD3612 | NADP-dependent alcohol dehydrogenase, cinnamyl alcohol dehydrogenase | vanillin degradation |
| | ACIAD2015 | putative alcohol dehydrogenase | vanillin degradation |
| | ACIAD2929 | putative alcohol dehydrogenase | vanillin degradation |
| | ACIAD1743 | putative oxydoreductase protein | vanillin degradation |

| | | | |
|---|---|---|---|
| *tgnC* | ACIAD2542 | Putative aldehyde dehydrogenase | vanillin degradation |
| *alrA* | ACIAD3616 | aldehyde reductase | vanillin degradation |
| | ACIAD1950 | putative iron-containing alcohol dehydrogenase | vanillin degradation |
| | ACIAD3642 | Putative aldehyde dehydrogenase | vanillin degradation |
